# Adolescent alcohol consumption produces long term alterations in response inhibition and orbitofrontal-striatal activity in a sex-specific manner

**DOI:** 10.1101/2024.12.15.628593

**Authors:** Aqilah M McCane, Lo Kronheim, Bita Moghaddam

## Abstract

Alcohol use disorder (AUD) is strongly associated with initiation of drinking during adolescence. Little is known about neural mechanisms that produce the long-term detrimental effects of adolescent drinking. A critical feature of AUD is deficits in response inhibition, or the ability to withhold a reward-seeking response. Here we sought to determine if adolescent drinking affects response inhibition and encoding of neural events by the orbitofrontal cortex (OFC) and dorsomedial striatum (DMS), two regions critical for expression of response inhibition. Adolescent male and female rats were given access to alcohol for four hours a day for five consecutive days. Then, rats were tested in a cued response inhibition task as adolescents or adults while we recorded concomitantly from the OFC and DMS. Adolescent voluntary alcohol drinking impaired response inhibition and increased alcohol drinking in adult male but not female rats. Adolescent alcohol drinking also resulted in sex-specific effects on both unit firing and local field potential measures in the OFC and DMS during premature and correct actions. Collectively, these data suggest sex-specific effects of adolescent alcohol drinking on response inhibition and corresponding alterations in cortical-striatal circuitry.

**Highlights:**

- Moderate adolescent alcohol drinking disrupts adult response inhibition
- Action encoding in the OFC and DMS changes after adolescent alcohol drinking
- OFC-DMS connectivity is altered in males after adolescent alcohol drinking
- Adolescent alcohol drinking promotes increased alcohol intake in adult males

## INTRODUCTION

Exposure to alcohol during adolescence is associated with greater likelihood of developing an alcohol use disorder (AUD) in adulthood (Buchmann et al., 2009; Spear and Swartzwelder, 2014; Crews et al., 2016; Kyzar et al., 2016). Adults with AUD consistently display deficits in response inhibition, or the ability to withhold a reward-seeking response (Noel et al., 2007; Lawrence et al., 2009; Papachristou et al., 2013; Naim-Feil et al., 2014). Behavioral inhibition problems are also a risk factor for alcohol dependence (Rubio et al., 2008), and can be used to differentiate populations at risk for alcohol-related problems (Nigg et al., 2006). However, most of the mechanistic animal studies related to alcohol abuse and AUD have focused on adult models of alcohol use, leading to a fundamental paucity of mechanistic data and animal models related to the lasting impact of adolescent alcohol use.

Preclinical models are well positioned to experimentally assess the impact of alcohol exposure on behavior in adolescence but are challenging to implement due to the short duration of rodent adolescence. In the current work, we utilized the Cued Response Inhibition Task (CRIT), which was designed to measure response inhibition in both adolescents and adults (Simon et al., 2013). Behavioral measurements of response inhibition were coupled with *in vivo* electrophysiology recordings in the lateral portion of the orbitofrontal cortex (OFC) and dorsomedial striatum (DMS). These regions were selected because the frontal cortex and striatum are amongst the last brain regions to mature and are therefore incredibly vulnerable during adolescence (Huttenlocher, 1979; Teicher et al., 1995; Lebel et al., 2008). Frontal-striatal connectivity during response inhibition is negatively correlated with severity of alcohol dependence (Courtney et al., 2013), further linking aberrations in cortico-striatal networks to AUD pathology. The OFC and DMS are critical substrates for response inhibition and lesions of either the OFC or dorsal striatum promote impaired response inhibition, and increased impulsive choice (Chudasama et al., 2003; Rieger et al., 2003; Eagle et al., 2007; Mar et al., 2011). Optogenetic stimulation of the OFC-striatum pathway was able to reverse deficits in response inhibition in a rodent model of compulsive behavior (Burguière et al., 2013). Lastly, inhibition of OFC neurons that project to the DMS results in increased premature actions during CRIT performance (McCane et al., 2024).

Using CRIT, we examined the impact of adolescent ethanol (EtOH) drinking on response inhibition while performing *in vivo* electrophysiology recordings in the OFC and DMS in male and female rats. Electrophysiology and behavior were measured during both adolescence and adulthood to evaluate the immediate and long-term effects of EtOH exposure.

## MATERIAL AND METHODS

### Subjects

Male and female adolescent (postnatal day (PND) 28-46, male N=21, female N=20) and adult (PND 60+, male N=18 female N=23) Long Evans rats were bred in-house. All rats were initially pair housed under temperature and humidity-controlled conditions using a 12-h reverse light/dark cycle. All animals were singled housed on PND 28, coinciding with adolescent surgical implantation of electrodes. Adolescents (PND 28) and adults (PND 60) were surgically implanted with custom-made 8-channel electrode arrays (50-µm-diameter tungsten wire insulated with polymide, California Fine Wire) in the OFC (AP 3.2, ML 3.0, DV-4.0) and DMS (AP 0.7, ML 1.6,DV -4.0) under isoflurane anesthesia as described previously (McCane et al., 2024). Behavioral testing occurred following one week of surgical recovery time. All experiments were performed during the dark phase, in accordance with the National Institute of Health’s Guide to the Care and Use of Laboratory Animals, and were approved by the Oregon Health and Science University Institutional Animal Care and Use Committee. After the end of each experiment, animals were anesthetized, perfused with paraformaldehyde, and histology was performed to confirm probe placements (Supplemental Figure 1). Animals with probes outside of the target regions were excluded from analyses.

### Statistical analyses

All analyses were performed in MATLAB (MathWorks) and R (https://www.r-project.org/). Unless otherwise specified, all comparisons were first tested using analyses of variance (ANOVA) testing, with between-subjects factors sex, age, and reward. All main effects were followed by post hoc tests for multiple comparisons.

### Alcohol exposure paradigms

A modified rat drinking-in-the-dark model (Holgate et al., 2017) was used to expose animals to EtOH. One hour into the dark (active) cycle, water bottles were replaced with two bottles, one containing water and the other containing either a 5% sucrose solution, or 20% v/v EtOH mixed in 5% sucrose and water. Bottles were fitted with double ball bearings to minimize leakage. Animals had access to solutions for four hours over the course of five days (PND 30-34). Animals were weighed prior to solution access so that intake in g/kg could be computed for each animal 30 minutes into drinking and at the end of the four-hour session.

After behavioral recordings and during adulthood (PND 80-93), all animals experienced a second home-cage drinking paradigm. An intermittent two-bottle choice paradigm was utilized to measure adult EtOH drinking behaviors (Simms et al., 2008). Similar to adolescent drinking, animals had two bottle access one hour into the dark cycle. However, unlike adolescent drinking, all animals had access to 20% v/v EtOH mixed in 5% sucrose and water. Two-bottle choice occurred every other day for 14 days, resulting in seven days of drinking. Intakes were measured 30 minutes after the two bottles were administered and 24 hours, at the cessation of the session.

### Behavior

Behavior occurred in an operant chamber (Coulbourn, Instruments) equipped with a food trough and reward magazine opposite a nose-poke port with a cue light, infrared photo-detector unit, and a tone-generating speaker. Adolescents and adults were food restricted, habituated to the operant box and trained to nose poke in response to a light cue for a sucrose pellet (45 mg, Bio-Serv) on a fixed ratio one schedule over two days. After successful acquisition of cue-action responding, all animals begin CRIT training as described previously (Simon et al., 2013; McCane et al., 2024). Each CRIT session lasted 60 minutes. Each trial began with the simultaneous presentation of two stimuli, an inhibitory cue (auditory tone) and an action cue (stimulus light). Responses made during the inhibitory cue were not rewarded and coded as “premature.” Following cessation of the inhibitory cue (variable time 5-30 s), the action cue remained illuminated for 10 seconds. An action executed during this time was rewarded and coded as “correct”. Failure to respond within 10 seconds was coded as an omission. Following either a response or omission, a variable length inter-trial interval of 10-12 seconds preceded the start of a new trial (Figure 1). Behavior data were analyzed using mixed design ANOVA testing with between subjects factors age, sex, reward group (whether rats received EtOH or sucrose as adolescents) and within subjects factor session. To understand group effects, data were aggregated across sessions 7-10 and the influence of age, sex and reward group was assessed using between subjects ANOVA testing and Tukey honest significance difference (HSD) post hoc testing, when appropriate.

**Figure 1.**
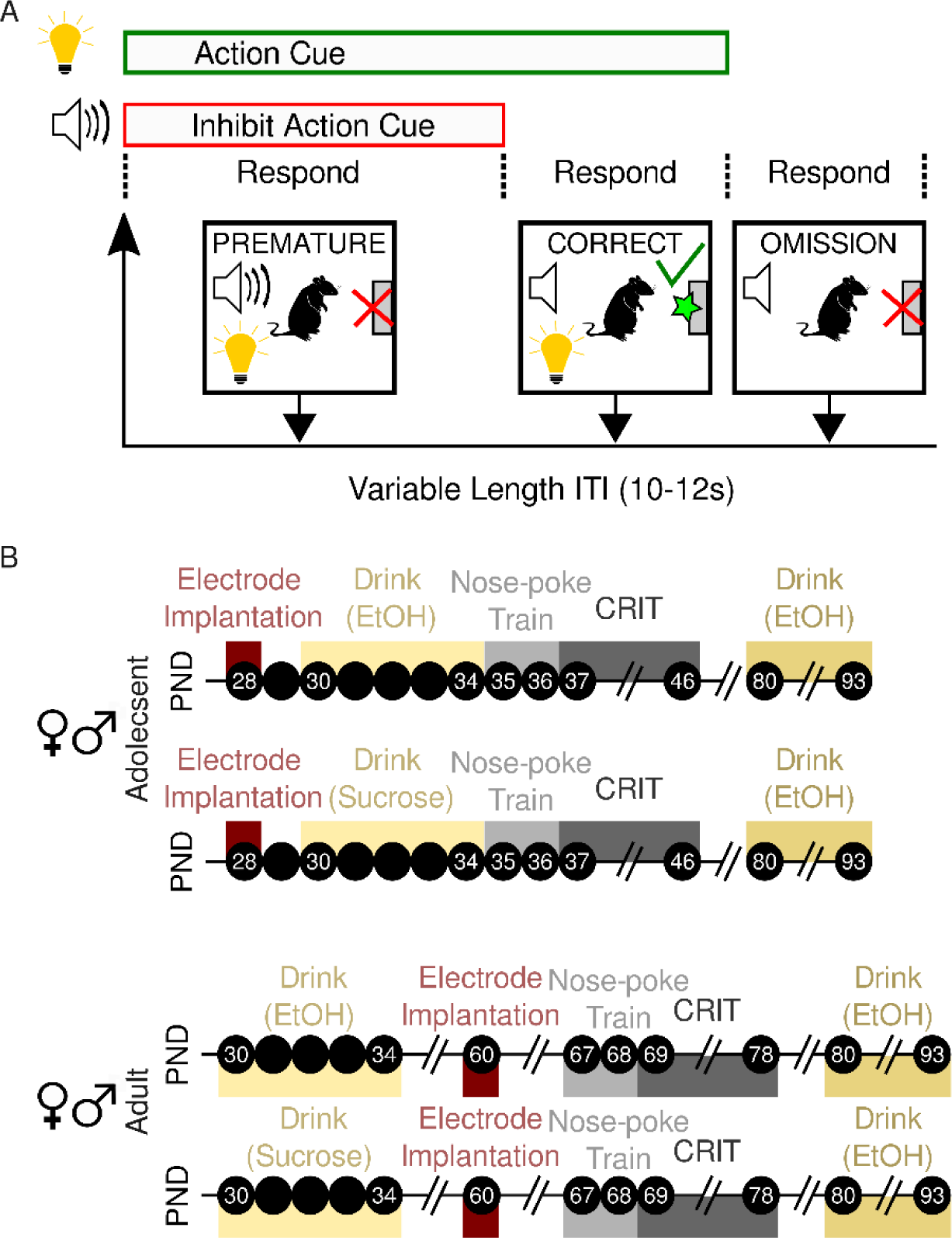
Schematic of experimental design. (A)The Cued Response Inhibition task (CRIT) yielded three response types. Responses made during the inhibit action cue were coded as “premature”, those made during the action cue, but in the absence of the inhibit action cue were coded as “correct”, and lack of response during either cue was coded as an “omission.” (B) Timeline depicting the postnatal days (PND; black circles) that experimental events occurred in male and female adolescent and adult rats.

### Electrophysiology Recordings

Single units and local field potentials (LFPS) were simultaneously recorded during performance of CRIT over the course of 10 sessions using a Plexon recording system. Spikes were amplified at 1000 X gain, digitalized at 40kHz, and single-unit data were band pass filtered at 300Hz. Single units were isolated using Kilosort (Allen et al., 2018). OFC neurons with baseline firing rates less than or equal to 10 Hz and spike widths greater than or equal to 0.30 were classified as putative pyramidal cells (Quirk et al., 2009), while DMS neurons with baseline firing rates less than or equal to 5 Hz and spike widths greater than or equal to 0.25 were classified as putative medium spiny neurons (Cayzac et al., 2011).

### Firing Rate analyses

Individual firing rates were first visualized by generating a heatmap of each unit’s firing rate, Z-score normalized to that unit’s baseline firing rate, the inter-trial interval (ITI) prior to cue onset. The percentage of units activated, inhibited or unchanged was computed by examining a 500 ms epoch following events of interest (tone on or action), measuring the mode Z-score for each unit, and subsequently calculating the number of units whose mode were greater than zero (activated), less than zero (inhibited) or equal to zero (unchanged). Chi-square test was used to determine if the proportion of activated, inhibited or unchanged units differed between reward groups, with at an alpha level of 0.05. Next, firing rates for all units were averaged across trials. To determine whether firing rate changed during events of interest, data were stratified by age and sex, and Friedman’s ANOVA was computed for each reward group. Significant main effects were followed up with Tukey’s HSD multiple comparison correction. Differences between reward groups were assessed using permutation tests (Jean-Richard-dit-Bressel et al., 2020).

### Spectral analyses

The Matlab wrapper for the fitting oscillation & one over f (FOOOF) toolkit was used to perform spectral analyses on LFP recordings (Donoghue et al., 2020). The aperiodic exponent or slope of the power spectral density was measured using a broad frequency range (1-50 Hz) as recommended (Donoghue et al., 2020; Ostlund et al., 2021). Welch’s power spectral density estimate was used to compute a power spectrum density, and putative theta oscillation power was extracted. Neither slope, nor power was computed from data sets which did not exhibit putative theta oscillations, as indicated by a frequency specific peak. Firing rate analyses indicated that action epochs were most strongly influenced by reward group. LFP analyses were therefore centered around correct and premature actions. Data were segregated by sex, consistent with previous analyses. The impact of reward on power and slope in each age group was assessed using ANOVA testing. Significant interactions were followed up using Tukey HSD post doc testing.

### Phase synchrony

Data were filtered in the theta band (5-11 Hz) and segregated by behavioral event. OFC-DMS synchrony was measured using the phase locking index γ which was computed by taking the complex value of the average of all points (1/N) where φ_1 (t) and φ_2 (t) are two phases from the filtered signals, the phase difference θ(t_j )=φ_1 (t_j )-φ_2 (t_j ), t_j are the times of data points, and N is the number of all data points during the given time interval (Lachaux et al., 1999; Hurtado et al., 2004; McCane et al., 2018).

### Regression analyses

Regression analyses were next employed to understand whether activity in the OFC and DMS differentially contributed to expression of behavior in EtOH or sucrose animals. Because activity around premature actions and in males was most influenced by adolescent EtOH exposure, these analyses investigated the relationship between premature action physiology and behavior in male rats only. The regression model was implemented using the MATLAB function *fitrensemble*. The following predictor variables were used: OFC power, OFC slope, OFC-DMS synchrony, DMS slope and DMS power. Predictor importance of all model variables was computed by summing the regression model estimates over all weak learners in the ensemble.

## RESULTS

### Adolescent male and female rats voluntarily consume EtOH in their home-cage

EtOH intake changed across days at both the 30 min (main effect of day: F(4,180)=3.33,p=0.01; Figure 2A) and the 4-hr timepoints (main effect of day: F(4,181)=7.67,p=9.86x10^-6^; Figure 2B). Sucrose intake increased across days at both the 30 min (main effect of day: F(4,171)=8.79,p=1.79x10^-6^; Figure 2C) and 4-hr timepoints (main effect of day: F(4,181)=7.67,p=9.86x10^-6^; Figure 2D). Overall males and females consumed similar amounts of EtOH and sucrose (p values >0.05), although there was a non-significant trend towards a main effect of sex during at the 4-hr timepoint in the sucrose animals (p=0.05).

**Figure 2.**
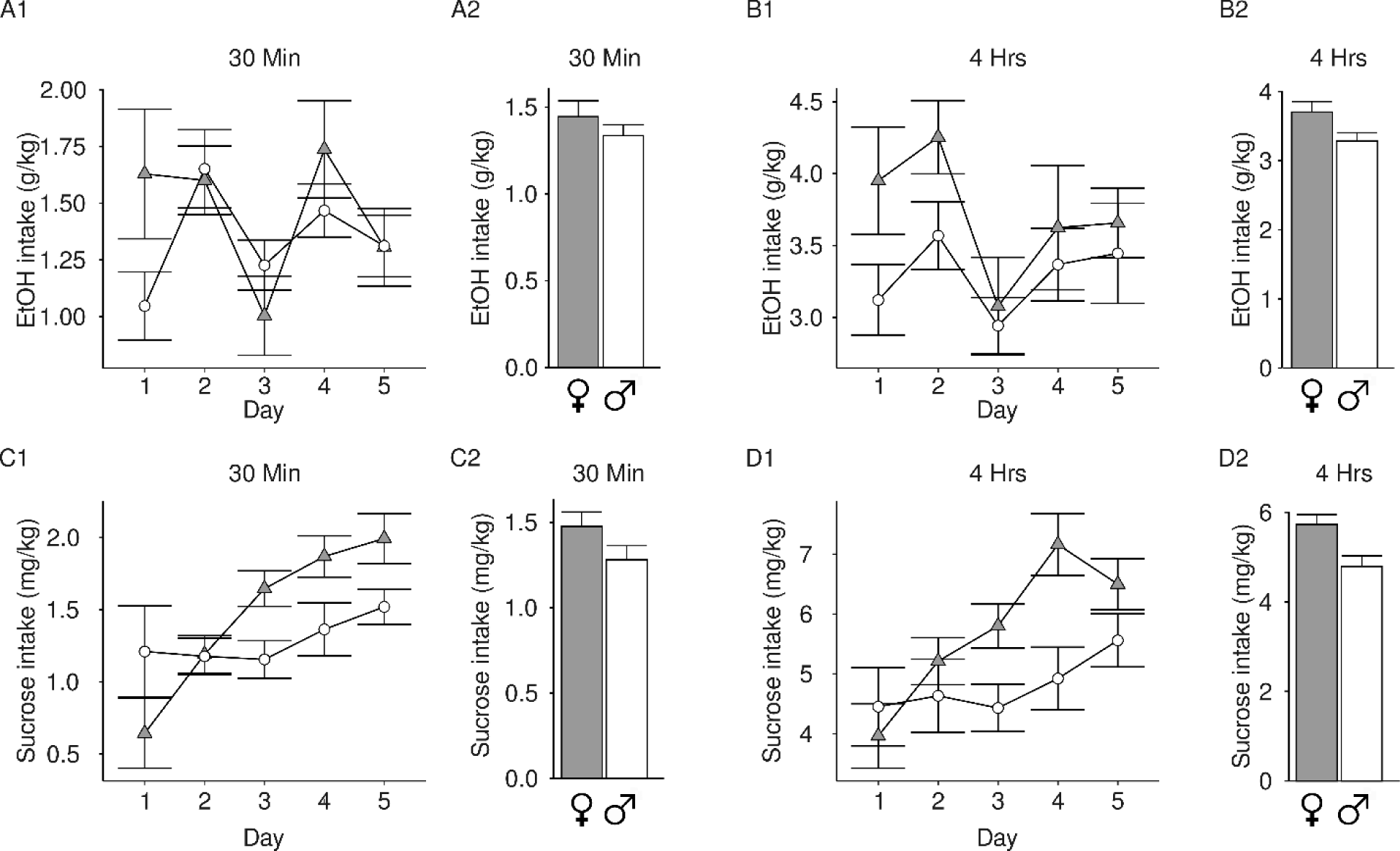
Adolescent home-cage drinking of EtOH or sucrose vehicle. (A-B) 20% v/v EtOH mixed in 5% sucrose or (C-D) 5% sucrose consumption in female (grey/triangle) or male (white/circle) adolescent rats was measured 30 min (1) or 4 hrs (2) after bottle placement in the cage. Bar graphs represent average of drinking across all days. A significant main effect of day was observed in all groups. (A1) Rats consumed varying amounts of EtOH intake after 30 min across days and (A2) both females and males consumed more than 1.0 g/kg within 30 min, on average. (B1) EtOH intake after 4 hrs of access changed across days. (B2) Females and males consume similar amounts of EtOH overall. (C1) Sucrose intake after 30 min of access increased over days. (C2) Females and males consume similar amounts of sucrose overall. (D1) Sucrose intake after 4 hrs of access increases over days and (D2) females and males consume similar amounts of sucrose overall. Data are presented as mean + SEM.

### Adolescent EtOH drinking produces sex-specific deficits in response inhibition

To understand the impact of adolescent EtOH drinking on CRIT performance, data were first divided by sex and age and analyzed using repeated measures ANOVA. Next, data were collapsed across the last four sessions and the influence of sex, drinking group and age on behavioral performance was investigated using between subject’s ANOVA with factors sex, reward and age.

Premature responses, in which rats failed to withhold reward-seeking actions during the inhibitory cue, increased over days for all groups (main effect of session: female adolescent: F(9,142)=5.07,p=6.22x10^-6^, female adult: F(9,195)=16.38,p<2x10^-16^, male adolescent: F(9,127)=3.27,p=0.001, male adult: F(9,146)=2.34,p=0.02; Supplemental Figure 2A). In the aggregated data, a significant reward × sex interaction for premature responses was observed (F(1,279)=11.89,p=0.0007; Figure 3A). Premature response data was therefore stratified by sex. Adolescent females made more premature responses than adult females (main effect of age: F(1,159)=22.50,p=4.65x10^-6^). Sucrose females also made more premature responses than EtOH females (main effect of reward: F(1,159)=6.38,p=0.001). In males, adolescents made more premature responses than adults (main effect of age F(1,120)=11.76,p=0.0008). Compared to sucrose males, EtOH males made more premature responses (main effect of reward: F(1,120)=11.88,p=0.0008), an effect driven by differences between sucrose and EtOH adults (Tukey post hoc, p=0.04).

**Figure 3.**
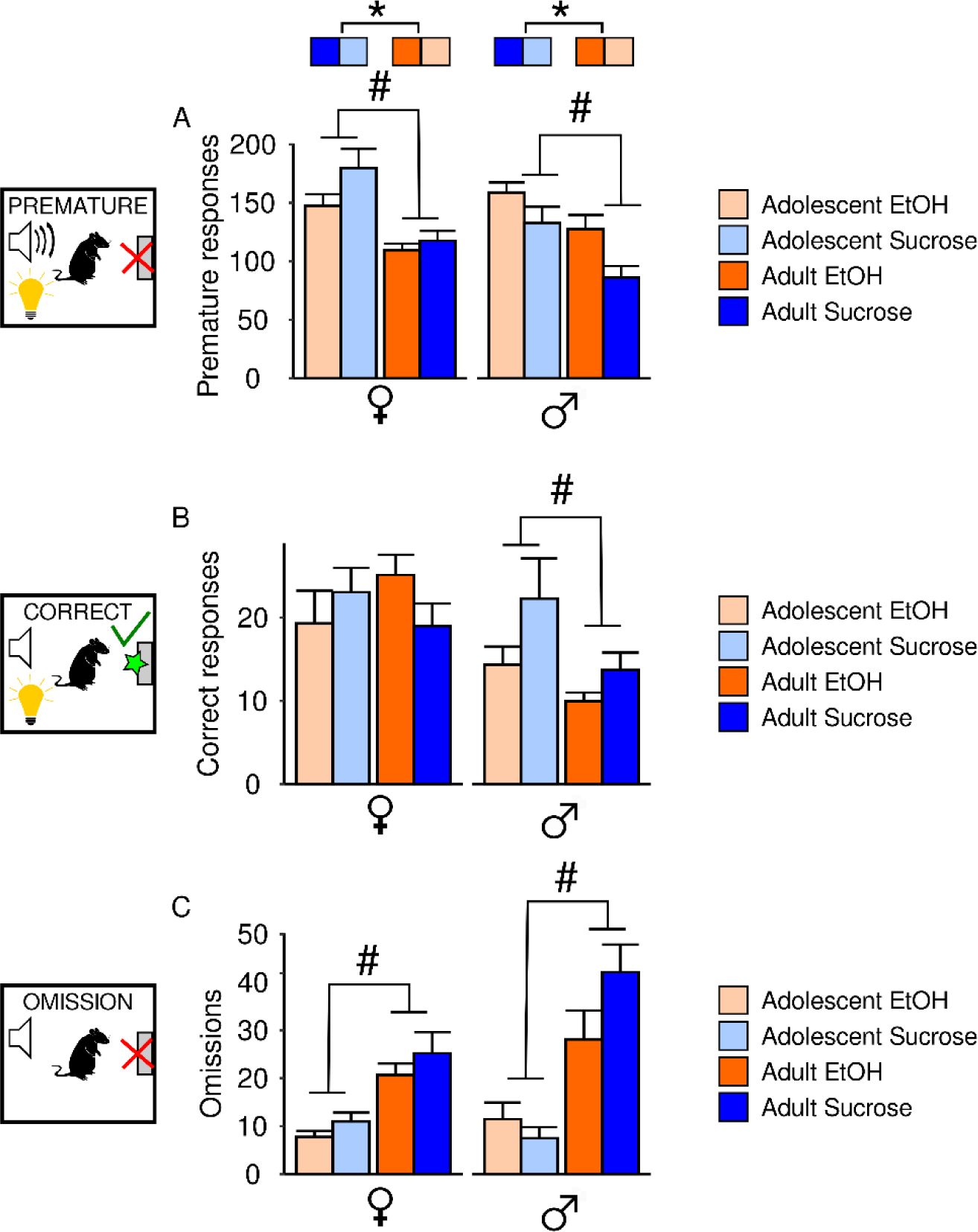
Impact of adolescent EtOH drinking on CRIT behavioral performance of male and female adolescents and adults. Three response types were assessed in CRIT: (A) Premature, (B) Correct, and (C) Omission. Male and female rats who drank EtOH (orange) or the vehicle sucrose (blue) performed ten days of CRIT. Lighter shades of each color reflect adolescents and darker shades reflect adults. Bar graphs depict behavioral performance for each group aggregated across the last four sessions of CRIT. (A) In females, adolescents make more premature responses than adults, and EtOH adolescents make fewer premature responses than the sucrose group. In males, adolescents make more premature actions than adults, and EtOH males make more premature actions than sucrose males. (B) Adolescent males make more correct responses than adult males. No significant effect of EtOH drinking was observed. (C) Adolescents of both sexes make fewer omissions compared to adults. No significant of EtOH drinking was observed. Data are presented as mean ± SEM. #p<0.05 main effect of age, *p<0.05 main effect of group.

In trials in which the animals correctly inhibited responding, the number of correct responses increased across sessions for all groups (main effect of session: female adolescent: F(9,142)=5.82,p=7.08x10^-7^, female adult: F(9,195)=17.0,p<2x10^-16^, male adolescent: F(9,127)=4.43,p=4.72x10^-5^, male adult: F(9,146)=2.78,p=0.005; Supplemental Figure 2B). Correct responding was also stratified by sex following a significant main effect of sex (F(1,279)=14.54,p=0.0002; Figure 3B). However, neither reward nor age influenced female correct responding (p values >0.50). In contrast, males in the alcohol group had an apparent, but not significant decrease in correct responding (F(1,120)=3.70,p=0.06), and age significantly influenced the number of correct responses male rats made (main effect of age: F(1,120)=6.20,p=0.14).

All groups decreased the number of trials omitted across sessions: (main effect of session: female adolescent: F(9,142)=11.13,p=4.92x10^-13^, female adult: F(9,195)=34.38,p<2x10^-16^, male adolescent: F(9,127)=5.63,p=1.54x10^-6^, male adult: F(9,146)=12.14,p=3.4x10^-14^; Supplemental Figure 2C). Overall, fewer omission trials were observed in adolescents compared to adults (main effect of age :F(1,279)=45.53,p=8.66x10^-11^) and females compared to males (main effect of sex: F(1,279)=7.40,p=0.007; Figure 3C).

Beyond the three trial outcomes, the response inhibition ratio (RIR), latency to make premature responses and number of nose pokes and trough entries during the ITI were assessed (Supplemental Figure 3). RIR was computed by taking the number of correct trials over the number of premature trials for each animal as reported previously (Simon et al., 2013).

The RIR increased over sessions in female adolescents (main effect of session: F(9,124)=2.37,p=0.02), and adults (main effect of session: F(9,195)=5.01,p=4.42x10^-6^) but not in males, (p values >0.05). In the aggregated data, a significant interaction between reward and sex was observed (F(1,279)=6.33,p=0.01). Data were therefore stratified by sex, which yielded no significant main effects in females (all p values> 0.05) but a main effect of reward in males (F(1,120)=5.72,p=0.02), indicating that males who drank EtOH in adolescence had a smaller RIR.

Latency to make a premature response changed across sessions in female adolescents (main effect of session: F(9,139)=9.74,p=1.84x10^-11^), female adults (main effect of session: F(9,193)=2.04,p=0.04), and male adults (main effect of session: F(9,135)=2.180,p=0.0271). When the last 4 sessions were aggregated, a significant main effect of sex (F(1,275)=4.90,p=0.03), as well as a sex by reward interaction was observed (F(1,275)=10.62,p=0.001). When data were subsequently stratified by sex, latency to make a premature response was unaffected by reward or age in females (all p values >0.05). In contrast, males who were exposed to EtOH in adolescence made premature actions sooner than sucrose males (main effect of reward: F(1,116)=11.55,p=0.0009).

Lastly, behavior during the ITI was analyzed. Number of nose-pokes during the ITI changed across session in female adolescents (main effect of session: F(9,139)=6.99,p=2.77x10^-8^) and female adults (main effect of session: F(9,191)=2.82, p=0.004), but not males (p values >0.05). Females and males made different numbers of responses during the ITI (main effect of sex: F(1,272)=11.33,p=0.0009), promoting a stratification of data by sex. In females, reward and age did not influence responding during the ITI (all p values >0.05), but adolescent EtOH use and age both influenced ITI nose pokes in males (F(1,114)=8.27,p=0.005). Post hoc testing indicated that adult EtOH males made more ITI nose pokes than sucrose males (Tukey post hoc, p=0.0001). Number of trough entries during the ITI changed across session in female adults (main effect of session: F(9,191)=3.53,p=0.0005), but no other group (p values >0.05). In the aggregated data, only age influenced the number of reward trough entries during the ITI (main effect of age: F(1,275)=12.78,p=0.0004).

### Adolescent EtOH drinking produces sex-specific alterations of premature action encoding

Because sex differences in CRIT behavior were observed, the impact of adolescent EtOH drinking on single unit activity was analyzed in male and female rats. Table 1 depicts statistical test results of ANOVA testing for all groups.

**Table 1.** Friedman ANOVA results.

| OFC |  |  |  |  |  |  |
| --- | --- | --- | --- | --- | --- | --- |
|  |  | Premature Cue | Premature Action | Correct Cue | Correct Action | Omission Cue |
| Adolescent | Female Sucrose | $\chi^2(39)=70.87$,<br>$p=0.001$ | $\chi^2(39)=131.03$,<br>$p=7.04 \times 10^{-12}$ | $\chi^2(39)=49.14$,<br>$p=0.13$ | $\chi^2(39)=70.51$,<br>$p=0.002$ | $\chi^2(39)=131.78$,<br>$p=5.37 \times 10^{-12}$ |
| | Female EtOH | $\chi^2(39)=66.06$,<br>$p=0.004$ | $\chi^2(39)=141.62$,<br>$p=1.45 \times 10^{-13}$ | $\chi^2(39)=59.12$,<br>$p=0.02$ | $\chi^2(39)=116.75$,<br>$p=1.10 \times 10^{-9}$ | $\chi^2(39)=68.54$,<br>$p=0.002$ |
| | Male Sucrose | $\chi^2(39)=47.56$,<br>$p=0.1635$ | $\chi^2(39)=160.17$,<br>$p=1.27 \times 10^{-16}$ | $\chi^2(39)=65.98$,<br>$p=0.004$ | $\chi^2(39)=95.23$,<br>$p=1.32 \times 10^{-6}$ | $\chi^2(39)=56.13$,<br>$p=0.04$ |
| | Male EtOH | $\chi^2(39)=190.98$,<br>$p=6.41 \times 10^{-22}$ | $\chi^2(39)=697.81$,<br>$p=1.24 \times 10^{-121}$ | $\chi^2(39)=100.1$,<br>$p=2.83 \times 10^{-7}$ | $\chi^2(39)=90.93$,<br>$p=4.97 \times 10^{-6}$ | $\chi^2(39)=96.54$,<br>$p=8.77 \times 10^{-7}$ |
| Adult | Female Sucrose | $\chi^2(39)=72.15$,<br>$p=0.001$ | $\chi^2(39)=293.11$,<br>$p=1.09 \times 10^{-40}$ | $\chi^2(39)=71.36$,<br>$p=0.001$ | $\chi^2(39)=59.8$,<br>$p=0.02$ | $\chi^2(39)=53.98$,<br>$p=0.06$ |
| | Female EtOH | $\chi^2(39)=68.61$,<br>$p=0.002$ | $\chi^2(39)=748.44$,<br>$p=4.57 \times 10^{-132}$ | $\chi^2(39)=83.63$,<br>$p=4.29 \times 10^{-5}$ | $\chi^2(39)=138.57$,<br>$p=4.47 \times 10^{-13}$ | $\chi^2(39)=59.26$,<br>$p=0.02$ |
| | Male Sucrose | $\chi^2(39)=69.46$,<br>$p=0.002$ | $\chi^2(39)=203.45$,<br>$p=3.99 \times 10^{-24}$ | $\chi^2(39)=40.1$,<br>$p=0.42$ | $\chi^2(39)=25.42$,<br>$p=0.95$ | $\chi^2(39)=250.13$,<br>$p=1.28 \times 10^{-32}$ |
| | Male EtOH | $\chi^2(39)=37.79$,<br>$p=0.52$ | $\chi^2(39)=119.07$,<br>$p=4.92 \times 10^{-10}$ | $\chi^2(39)=55.72$,<br>$p=0.04$ | $\chi^2(39)=80.52$,<br>$p=0.0001$ | $\chi^2(39)=115.35$,<br>$p=1.78 \times 10^{-9}$ |
| DMS |  |  |  |  |  |  |
| Adolescent | Female Sucrose | $\chi^2(39)=42.12$,<br>$p=0.34$ | $\chi^2(39)=473.23$,<br>$p=5.66 \times 10^{-76}$ | $\chi^2(39)=52.52$,<br>$p=0.07$ | $\chi^2(39)=233.83$,<br>$p=1.29 \times 10^{-29}$ | $\chi^2(39)=45.65$,<br>$p=0.22$ |
| | Female EtOH | $\chi^2(39)=147.48$,<br>$p=1.61 \times 10^{-14}$ | $\chi^2(39)=562.09$,<br>$p=6.80 \times 10^{-94}$ | $\chi^2(39)=63.49$,<br>$p=0.008$ | $\chi^2(39)=275.89$,<br>$p=1.97 \times 10^{-37}$ | $\chi^2(39)=47.33$,<br>$p=0.17$ |
| | Male Sucrose | $\chi^2(39)=79.09$,<br>$p=0.0002$ | $\chi^2(39)=163.32$,<br>$p=3.75 \times 10^{-17}$ | $\chi^2(39)=56.95$,<br>$p=0.03$ | $\chi^2(39)=58.84$,<br>$p=0.02$ | $\chi^2(39)=54.64$,<br>$p=0.049$ |
| | Male EtOH | $\chi^2(39)=66.3$,<br>$p=0.004$ | $\chi^2(39)=274.09$,<br>$p=4.29 \times 10^{-37}$ | $\chi^2(39)=48.37$,<br>$p=0.144$ | $\chi^2(39)=121.85$,<br>$p=1.86 \times 10^{-10}$ | $\chi^2(39)=44.04$,<br>$p=0.27$ |
| Adult | Female Sucrose | $\chi^2(39)=130.29$,<br>$p=9.18 \times 10^{-12}$ | $\chi^2(39)=719.58$,<br>$p=4.09 \times 10^{-126}$ | $\chi^2(39)=50.13$,<br>$p=0.11$ | $\chi^2(39)=92.2$,<br>$p=3.38 \times 10^{-6}$ | $\chi^2(39)=67.43$,<br>$p=0.003$ |
| | Female EtOH | $\chi^2(39)=39.33$,<br>$p=0.46$ | $\chi^2(39)=214.03$,<br>$p=5.09 \times 10^{-26}$ | $\chi^2(39)=32.71$,<br>$p=0.75$ | $\chi^2(39)=161.59$,<br>$p=7.33 \times 10^{-17}$ | $\chi^2(39)=51.11$,<br>$p=0.09$ |
| | Male Sucrose | $\chi^2(39)=86.97$,<br>$p=1.63 \times 10^{-5}$ | $\chi^2(39)=240.1$,<br>$p=9.15 \times 10^{-31}$ | $\chi^2(39)=32.12$,<br>$p=0.77$ | $\chi^2(39)=118.22$,<br>$p=6.61 \times 10^{-10}$ | $\chi^2(39)=103.87$,<br>$p=8.36 \times 10^{-8}$ |
| | Male EtOH | $\chi^2(39)=98.41$,<br>$p=4.86 \times 10^{-7}$ | $\chi^2(39)=132.02$,<br>$p=4.92 \times 10^{-12}$ | $\chi^2(39)=66.14$,<br>$p=0.004$ | $\chi^2(39)=213.91$,<br>$p=5.37 \times 10^{-26}$ | $\chi^2(39)=65.43$,<br>$p=0.005$ |

In the OFC, adolescent EtOH drinking in females led to a larger increase in firing rate at the onset of the inhibitory cue and after premature actions compared to sucrose controls although the latter group had a greater proportion of units activated after premature actions (Figure 4A). In adults, premature actions were followed by a decrease in OFC firing rate relative to baseline in EtOH females which was absent in sucrose adults (Figure 4B). In the DMS, premature actions and the cues which preceded them were associated with an increase in firing rate in both EtOH and sucrose adolescent females (Figure 4C). Female adults who drank sucrose in adolescence exhibited an increase in DMS firing rate following cue presentation, likely driven by subpopulation of excited neurons (Figure 4D). DMS firing rate was suppressed following premature actions in both EtOH and sucrose adult females, but to a greater extent in sucrose rats (Figure 4D).

**Figure 4.**
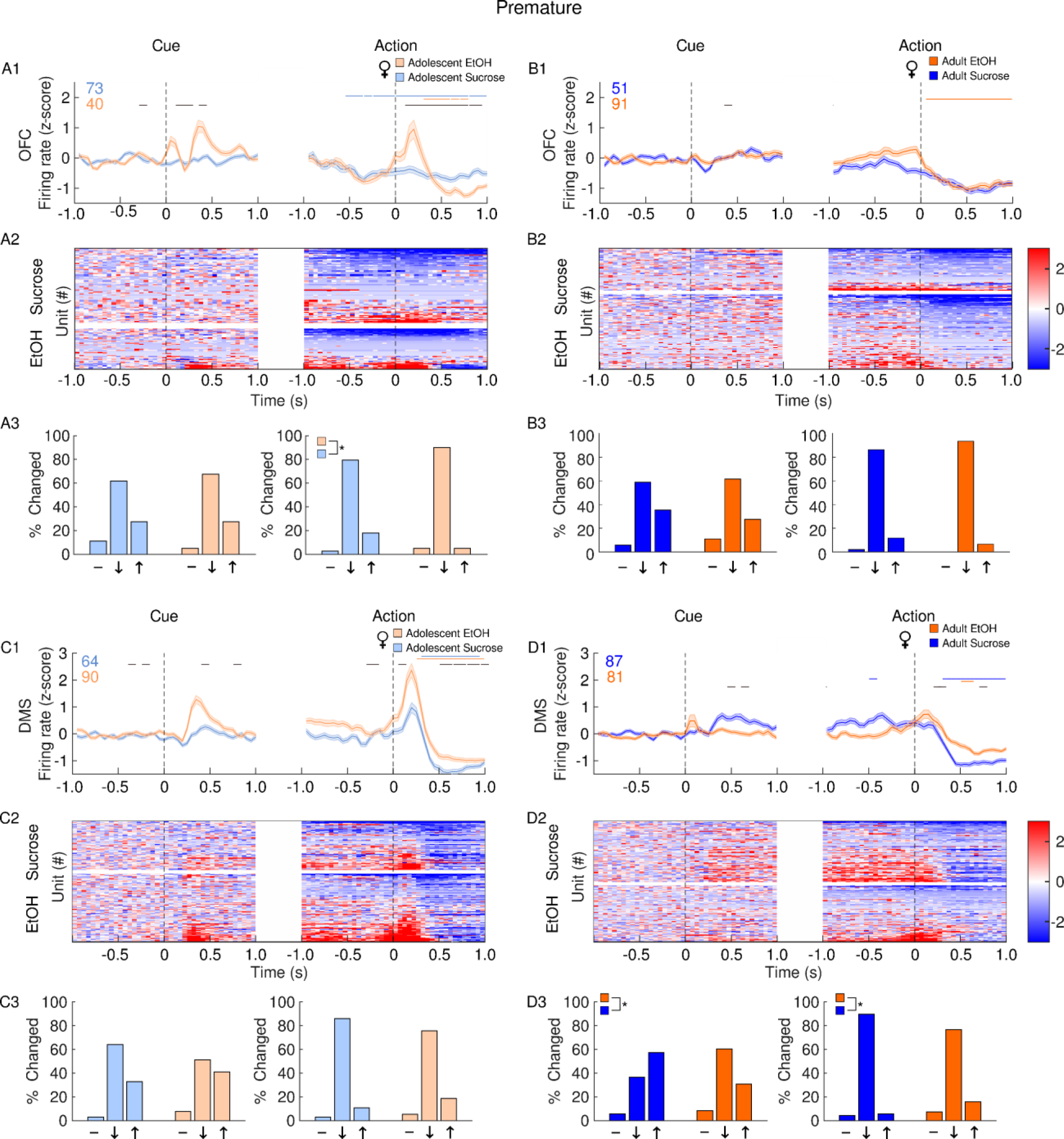
Impact of adolescent EtOH drinking on single unit activity in female adults and adolescents after premature actions during CRIT performance. Recordings were performed in the OFC (A-B) and DMS (C-D). Panels depict averaged firing rates (1), heatmaps of individual neural responses (2) and histograms (3) of the distribution of units unchanged (-), inhibited (↓) or excited (↑), 500 ms after events of interest (cue or action) female rats with a history of EtOH (orange) or sucrose (blue). Adolescents are plotted in the lighter shade of both colors. Events occur at time 0. (A1) Following premature responses, female EtOH adolescents show an increase in OFC firing rate which is absent in sucrose adolescents. (A2) Heatmaps of individual units show both sucrose and EtOH adolescents have a subpopulation of excited neurons prior to and after premature responses. (A3) Compared to sucrose rats, adolescent EtOH females show a greater proportion of neurons inhibited after premature actions. (B1) In adult females, premature responses are accompanied by a suppression in OFC firing rate in EtOH but not sucrose females. (B2) In both EtOH and sucrose animals, a population of neurons become inhibited following premature actions. (B3) EtOH and sucrose female adults have a similar distribution of neural responses following premature actions. (C1) During premature trials, EtOH adolescent females show a greater increase in DMS firing rate following cue presentation and premature actions compared to sucrose females. (C2) Both cue presentations and premature actions are followed by an increase in excitation in both EtOH and sucrose female adolescent rats. (C3) The distribution of neural responses is similar between EtOH and sucrose rats. (D1) Compared to EtOH rats, sucrose adult females show a larger increase in DMS firing rate following cue presentation and steeper decrease in firing rate following premature actions. (D2) In adult female sucrose rats, a subpopulation of neurons is excited following cue presentation and inhibited following premature actions. (D3) Compared to EtOH rats, sucrose rats have a greater proportion of cells excited following premature-trial cue presentation and a greater proportion of cells inhibited following premature actions. Data are presented as mean ± SEM. Colored numbers reflect the number of units recorded in each group. Colored significant bars represent Tukey HSD post hoc testing (p<0.05), compared to baseline firing rate for each group (EtOH or sucrose). Black significant bars reflect permutation testing between reward groups, p<0.05. *p<0.05 group difference, Chi-squared test

In both EtOH and sucrose control adolescent males, OFC firing rate was reduced during premature actions with each group exhibiting similar populations of neurons which became inhibited around premature actions (Figure 5A). In sucrose but not EtOH adult males, OFC activity was suppressed after premature actions (Figure 5B). In adolescent male rats, adolescent EtOH consumption was associated with an increase in DMS firing rate following premature actions, but the proportion of excited neurons was not statistically different from sucrose males (Figure 5C). In adult males, premature actions were followed by a larger increase in DMS firing rate in sucrose animals, but EtOH animals had a larger proportion of neurons excited following premature actions (Figure 5D).

**Figure 5.**
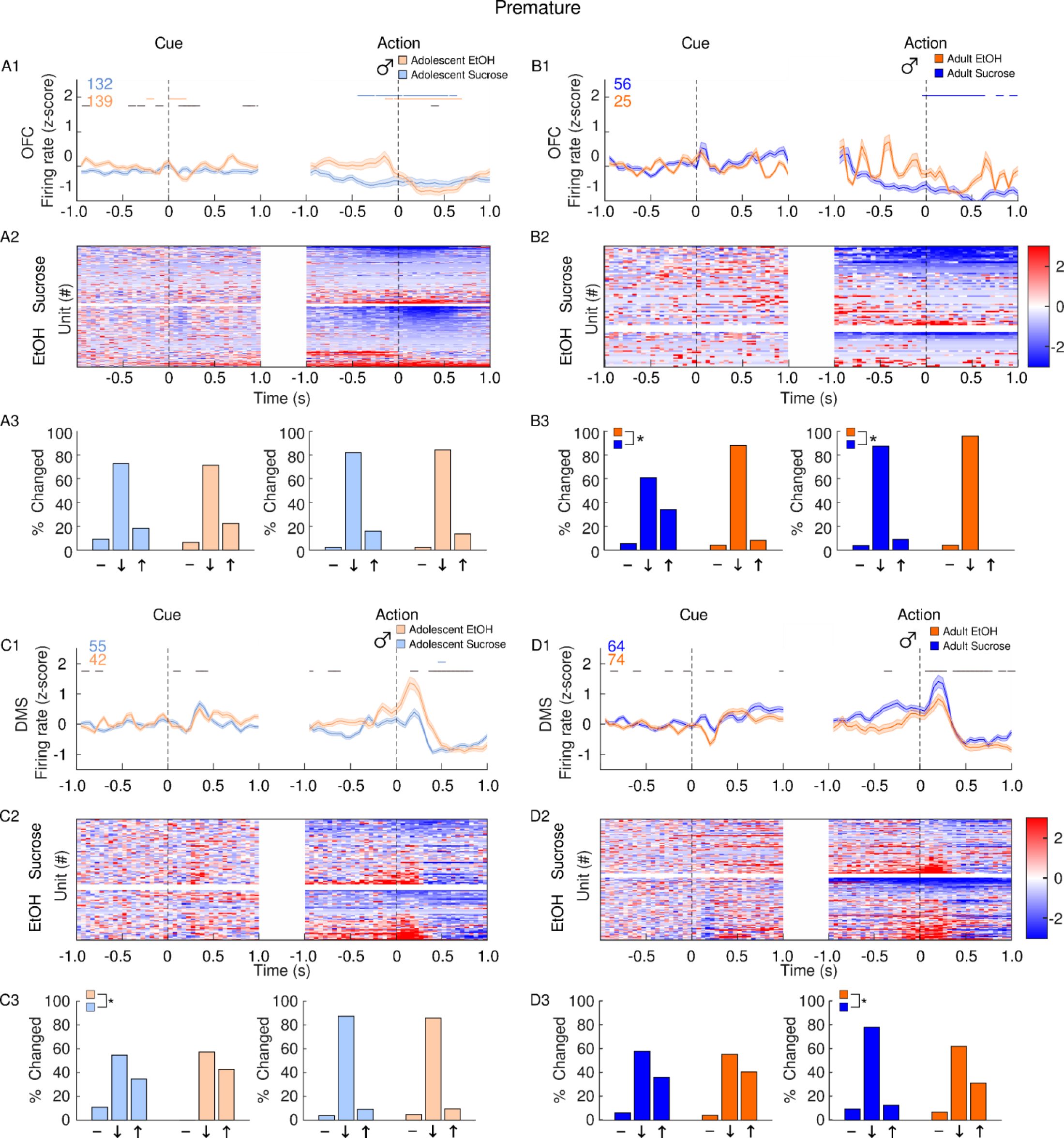
Impact of adolescent EtOH drinking on single unit activity in male adults and adolescents after premature actions during CRIT performance. Recordings were performed in the OFC (A-B) and DMS (C-D). Panels depict averaged firing rates (1), heatmaps of individual neural responses (2) and histograms (3) of the distribution of units unchanged (-), inhibited (↓) or excited (↑), 500 ms after events of interest (cue or action) male rats with a history of EtOH (orange) or sucrose (blue). Adolescents are plotted in the lighter shade of both colors. Events occur at time 0. (A1) Following premature responses, EtOH adolescent males show a suppression in OFC firing rate. (A2) Heatmaps show a subpopulation of neurons become suppressed following premature actions in EtOH adolescent males. (A3) EtOH and sucrose male adolescents have a similar distribution of neural responses following premature actions. (B1) EtOH male adult OFC firing rate following premature actions shows no clear pattern. (B2) Both EtOH and sucrose adult males exhibit a subpopulation of inhibited neurons surrounding premature actions. (B3) Following cue presentation and premature actions, EtOH adult males have a greater proportion of inhibited neurons. (C1) sucrose adolescent males show a larger increase in DMS firing rate following cue presentation, but EtOH adolescents show a larger response following premature actions. (C2) both EtOH and sucrose adolescent males show a subpopulation of neurons become excited following cue presentation and premature actions, but neurons excited following premature actions become inhibited within 0.5 ms in both groups. (C3) The distribution of neural responses differs between sucrose and adolescent males following cue presentation, but not following premature actions. (D1) Compared to sucrose rats, EtOH adult males show a greater suppression in DMS firing rate following cue presentation and an attenuated response following premature actions. (D2) a subpopulation of adult male EtOH neurons is inhibited following cue presentation, and both EtOH and sucrose rats have a population of cells excited following premature actions. (D3) Following premature actions, sucrose adult males have a greater proportion of inhibited neurons and a smaller proportion of excited neurons compared to EtOH rats. Data are presented as mean ± SEM. Colored numbers reflect the number of units recorded in each group. Colored significant bars represent Tukey HSD post hoc testing (p<0.05), compared to baseline firing rate for each group (EtOH or sucrose). Black significant bars reflect permutation testing between reward groups, p<0.05. *p<0.05 group difference, Chi-squared test

### Adolescent EtOH exposure produces sex-specific changes in correct action encoding

In the OFC, EtOH adolescent females exhibited a large increase in firing rate, driven by a small population of excited cells after correct actions (Figure 6A). OFC firing rate in adult EtOH females was increased following correct actions, but the proportion of cells excited by reward was similar between EtOH and sucrose rats (Figure 6B). In the DMS, a comparable increase in firing rate was observed after correct actions in both EtOH and sucrose female adolescents (Figure 6C) and adults (Figure 6D).

**Figure 6.**
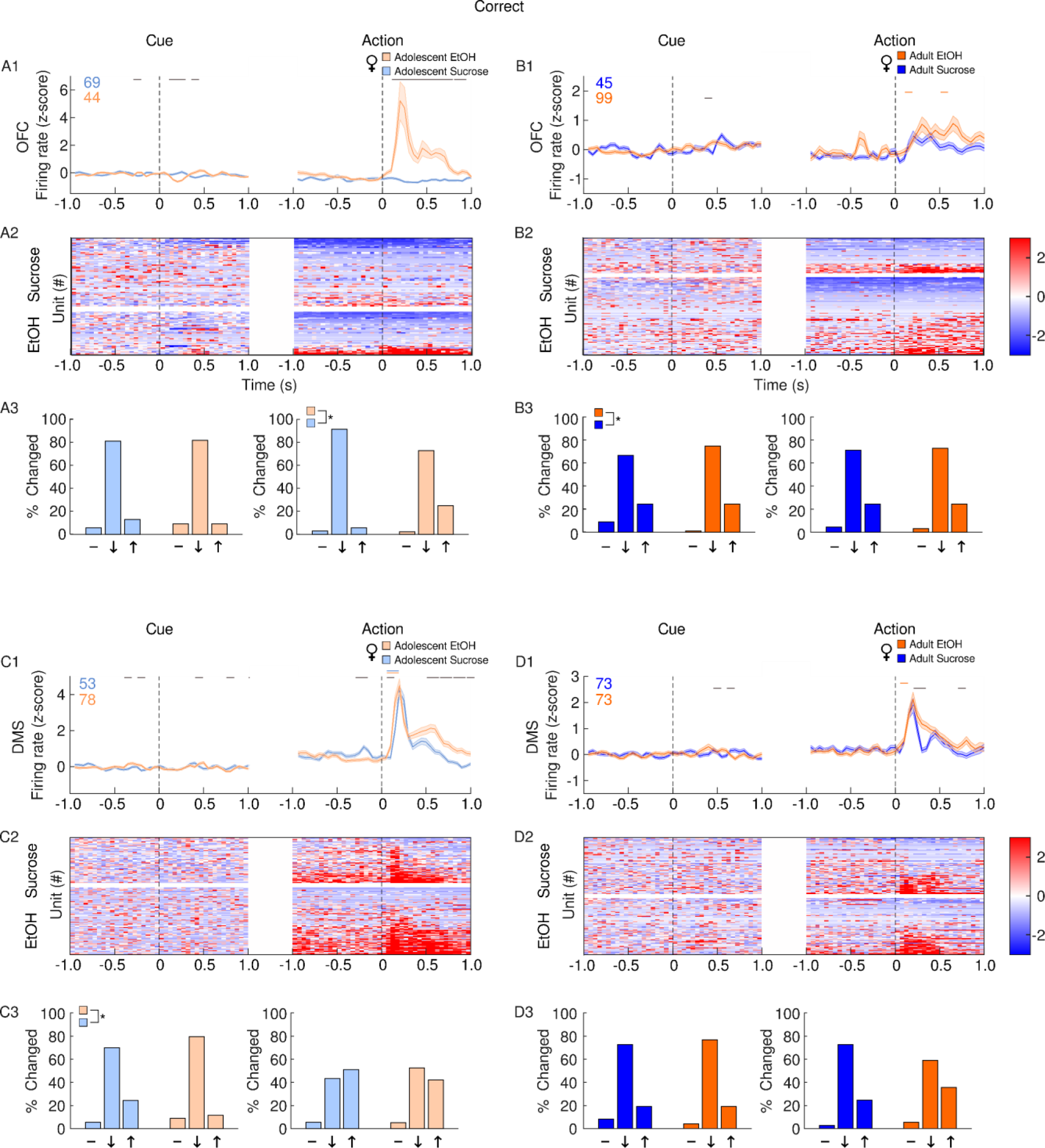
Impact of adolescent EtOH drinking on single unit activity in female adults and adolescents after correct actions during CRIT performance. Recordings were performed in the OFC (A-B) and DMS (C-D). Panels depict averaged firing rates (1), heatmaps of individual neural responses (2) and histograms (3) of the distribution of units unchanged (-), inhibited (↓) or excited (↑), 500 ms after events of interest (cue or action) male rats with a history of EtOH (orange) or sucrose (blue). Adolescents are plotted in the lighter shade of both colors. Events occur at time 0. (A1) Adolescent EtOH females show a larger increase in OFC firing rate following correct actions compared to sucrose females. (A2) Increased firing rate is observed in a subpopulation of EtOH, but not sucrose adolescent females. (A3) EtOH adolescent females have a greater proportion of excited cells after correct actions compared to sucrose rats. (B1) Adult EtOH females show a moderate increase in OFC firing rate following correct actions. (B2) Both Sucrose and EtOH female adults have a proportion of neurons excited following correct actions. (B3) The distribution of neural responses differ between reward groups in adult females after cue presentation, but not following correct actions. (C1) Both EtOH and sucrose adolescent females show an increase in DMS firing rate after correct actions. (C2) Cues which precede correct actions are followed by a larger proportion of excited neurons in sucrose adolescent females. (D1) Correct actions are followed by an increase in DMS firing rate in both sucrose EtOH exposed male adults. (D2) A subpopulation of DMS neurons are excited following correct actions in both sucrose and EtOH female adult rats. (D3) The distribution of neural responses following correct actions is similar between sucrose and EtOH female adults. Data are presented as mean ± SEM. Colored numbers reflect the number of units recorded in each group. Colored significant bars represent Tukey HSD post hoc testing (p<0.05), compared to baseline firing rate for each group (EtOH or sucrose). Black significant bars reflect permutation testing between reward groups, p<0.05. *p<0.05 group difference, Chi-squared test

During correct trials, EtOH adolescent male OFC activity was suppressed after cue presentation when compared to sucrose adolescent males, but OFC response to correct actions was overall similar between adolescent EtOH and sucrose males (Figure 7A). Similarly, adult males who drank EtOH in adolescence exhibited a greater percentage of inhibited OFC neurons after cue presentation during correct action trials (Figure 7B). Compared to adolescent sucrose males, adolescent EtOH males had a larger DMS response after correct actions and a corresponding increase in cells activated following correct actions (Figure 7C). In adults, sucrose males had a larger DMS response after correct actions and an apparent but not significant difference in the proportion of cells activated, compared to EtOH rats (Figure 7D).

**Figure 7.**
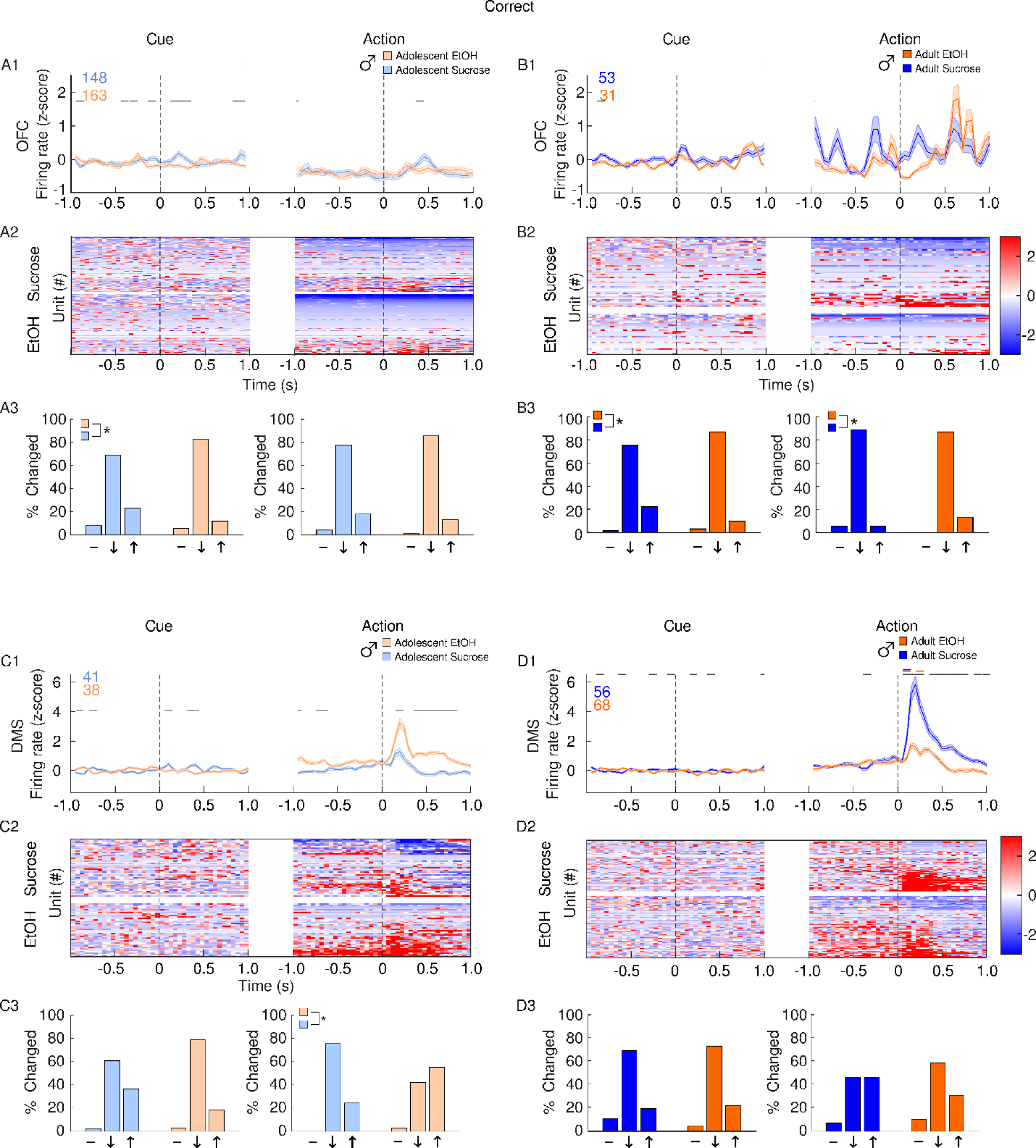
Impact of adolescent EtOH drinking on single unit activity in male adults and adolescents after correct actions during CRIT performance. Recordings were performed in the OFC (A-B) and DMS (C-D). Panels depict averaged firing rates (1), heatmaps of individual neural responses (2) and histograms (3) of the distribution of units unchanged (-), inhibited (↓) or excited (↑), 500 ms after events of interest (cue or action) male rats with a history of EtOH (orange) or sucrose (blue). Adolescents are plotted in the lighter shade of both colors. Events occur at time 0. (A1) Sucrose adolescent males exhibit a higher OFC firing rat following correct actions, and cues which preceded them. (A2) In adolescent sucrose males, a subpopulation of OFC neurons become more active following correct actions. (A3) Adolescent males in the sucrose group have a greater proportion of cells activated following correct trial cue presentation. (B1) In adults, OFC firing rates following both correct actions and cues which precede them do not differ from baseline activity. (B2) In adult sucrose exposed males, a subpopulation of OFC neurons become excited immediately following correct actions. (B3) A greater proportion of neurons are excited following correct trial cues in the sucrose group, while a greater proportion of EtOH neurons are excited following correct actions in the EtOH group. (C1) Compared to sucrose animals, EtOH exposed adolescent males have a larger increase in DMS firing rate following correct actions. (C2) In sucrose, but not EtOH adolescent males, a subpopulation of DMS neurons become inhibited following correct actions. (C3) Adolescent sucrose exposed male rats have a larger proportion of neurons inhibited following correct actions compared to EtOH exposed rats. (D1) Both EtOH and sucrose adult males show an increase in DMS firing rate following correct actions, but the sucrose response is larger. (D2) A large proportion of excited DMS neurons are observed in both EtOH and sucrose adult males following correct actions. (D3) The distribution of neural responses following correct actions is similar between sucrose and EtOH male adults. Data are presented as mean ± SEM. Colored numbers reflect the number of units recorded in each group. Colored significant bars represent Tukey HSD post hoc testing (p<0.05), compared to baseline firing rate for each group (EtOH or sucrose). Black significant bars reflect permutation testing between reward groups, p<0.05. *p<0.05 group difference, Chi-squared test

### EtOH exerts sex and age-specific effects on LFPs

The global impact of adolescent EtOH drinking was assessed by evaluating three LFP measures: the aperiodic component (slope), power and phase synchrony. These analyses focused on activity surrounding premature and correct actions only because adolescent EtOH drinking exerted subtle effects on cue presentation during omission trials (Supplemental Figure 4).

The aperiodic exponent was computed first by taking the slope of the power spectrum density (Figure 8A). Similar to behavioral and single unit data, data were divided by sex. In females, both age and reward group influenced OFC slope following premature actions (reward × age interaction: F(1,879)=7.06,p=0.008), where adults overall had lower exponents than adolescents, and adults who had adolescent EtOH experience exhibited a larger exponent compared to sucrose adults (Tukey post hoc, p values<0.05; Figure 8B). In males, OFC slope following premature actions was influenced by both reward and age (F(1,306)=16.04,p=7.78x10^-5^), where EtOH adolescents had a larger exponent compared to sucrose adolescents, and adolescents in both reward groups had lower exponents than adults (Tukey post hoc, p values <0.05; Figure 8B). In the DMS, adolescent alcohol exposure caused larger exponents in females after premature actions (main effect of reward: F(1,1116)=7.31,p=0.007; Figure 8C). In males, adults had a larger DMS exponent after premature actions (main effect of age: F(1,493)=97.74,p<2x10^-16^). While adolescent alcohol exposure differently influenced DMS slope for adults and adolescents (reward × age interaction: F(1,493)=8.20,p=0.004), reward differences did not survive post hoc testing (p values >0.05).

**Figure 8.**
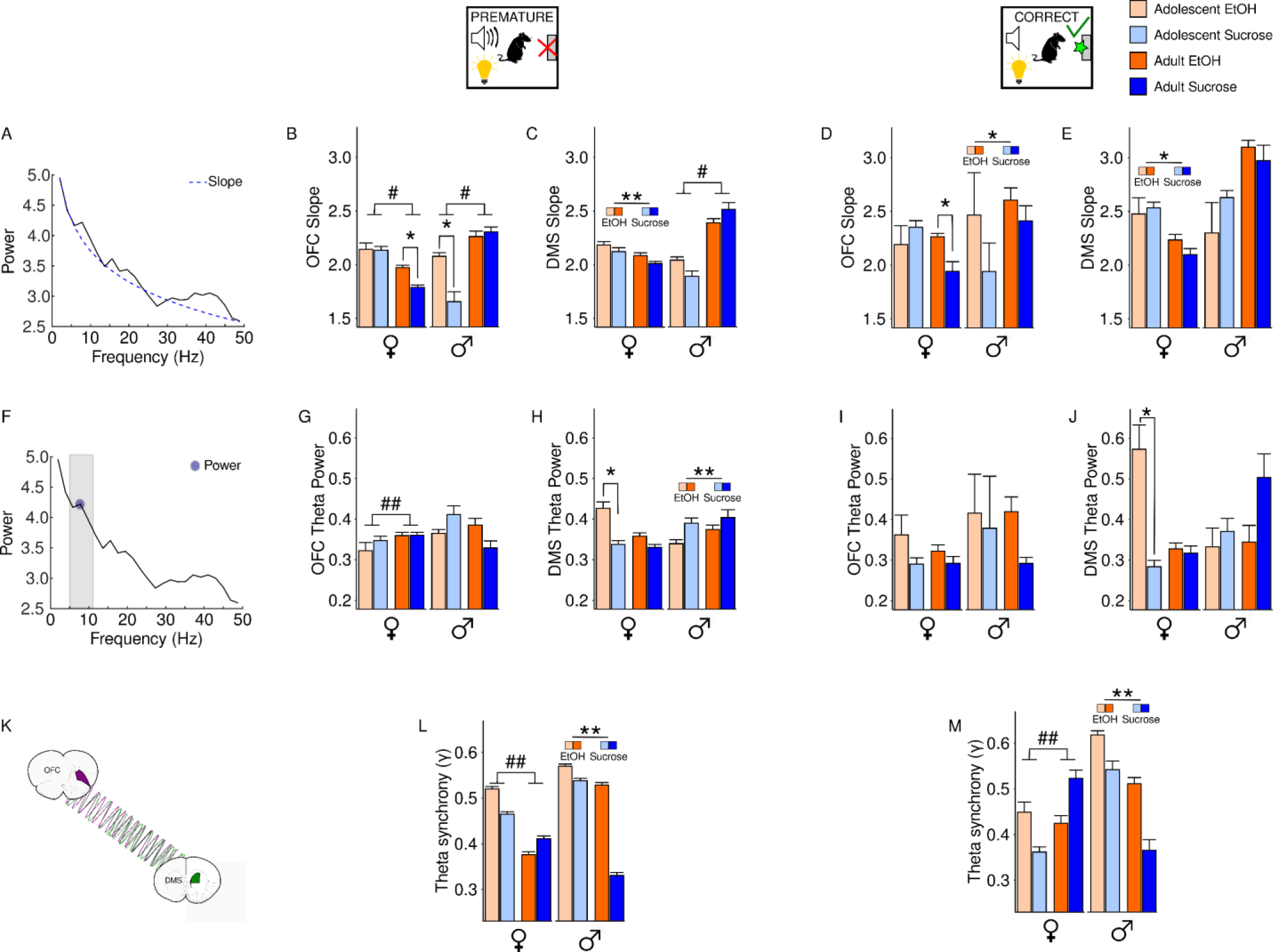
Local field potential analyses of OFC and DMS during CRIT performance in male and female adult and adolescents after adolescent EtOH drinking. Slope (A-E), Power (F-J) and OFC-DMS synchrony (K-M) were measured in male and female rats with a history of adolescent EtOH (orange) or sucrose (blue) exposure. (A) The aperiodic exponent was computed by taking the slope of the power spectrum density. (B) Following premature actions, female adults had lower OFC exponents than female adolescents, and female EtOH adults had a larger OFC exponent compared to female sucrose adults. In males, adolescents had lower OFC exponents compared to adults and EtOH adolescents had a larger OFC exponent compared to sucrose adolescents. (C) In females, adolescent alcohol exposures is associated with larger DMS exponents following premature actions. In males, DMS exponents were larger in adults compared to adolescents. (D) Following correct actions, female sucrose adolescents had a larger OFC exponent compared to female sucrose adults and EtOH adults exhibited a larger exponent compared to sucrose adults. In males, adolescent alcohol exposure was associated with larger OFC exponents following correct actions. (E) In the female group, DMS exponents were larger in adolescents compared to adults after correct actions. In contrast, male DMS exponents were not influenced by adolescent EtOH drinking. (F) Presence of a peak in the theta frequency range indicated the presence of theta oscillations which were extracted and used to measure theta power. (G) In females, adolescents exhibit reduced OFC theta power compared to adults following premature actions. In contrast, male adolescents exhibit stronger OFC theta power compared to adults. (H) Adolescent ETOH females had stronger DMS theta power compared to sucrose adolescents following premature actions. In males, ETOH was associated with decreased DMS theta power following premature actions. (I) OFC theta power following correct actions was not influenced by adolescent alcohol drinking in females or males. (J) following correct trials, ETOH adolescent females had stronger DMS theta power than sucrose adolescents, and adult males had stronger DMS theta compared to adolescents(K) OFC-DMS connectivity was measured by computing the phase-locking index gamma. (L) Following premature actions, adolescent females have stronger OFC-DMS synchrony compared to adult females while EtOH drinking in males was associated with increased OFC-DMS synchrony in both age groups. (M) Compared to adults, female adolescents exhibit weaker OFC-DMS synchrony following correct actions. Following correct actions, EtOH males exhibit stronger OFC-DMS connectivity compared to sucrose males. Data are presented as mean + SEM. ##p<0.05 main effect of age, **p<0.05 main effect of group, *p<0.05 Tukey post hoc group difference, #p<0.05 Tukey age difference.

After correct actions, a significant interaction between age and reward on OFC slope was observed in females (F(1,146)=8.41,p=0.004), where sucrose adolescents displayed a larger exponent compared to sucrose adults (Tukey post hoc, p=0.0002), and EtOH adults displayed a larger exponent compared to sucrose adults (Tukey post hoc, p=0.004; Figure 8D). In the DMS, adolescent alcohol exposure caused larger exponents in females after correct actions (F(1,173)=5.60,p=0.02), but had no effect in males (main effect of reward: F(1,46)=1.43,p=0.24; Figure 8E).

Next, theta power was computed by extracting the peak oscillation in the theta frequency range (Figure 8F). Following premature actions, female adolescents exhibited reduced OFC theta power compared to adults (main effect of age: F(1,879)=4.27,p=0.04), but there was no effect of reward group on theta power (main effect of reward: F(1,879)=0.10,p=0.75; Figure 8G). In males, a significant interaction between reward and age on OFC theta power after premature actions was observed (F(1,306)=8.10,p=0005), where sucrose-exposed adolescents had stronger theta power compared to sucrose adults (Tukey post hoc, p=0.02). Notably, there was a non-significant trend towards stronger theta power in EtOH adults compared to sucrose adults (p=0.08). In the DMS, a significant interaction between age and reward was observed in females after premature actions (reward × age interaction: F(1,1116)=8.07,p=0.005). Post hoc testing indicated that adolescent females with EtOH exposure had stronger DMS theta power than sucrose adolescents (Tukey post hoc, p= p=0.000002; Figure 8H). In males, adolescent alcohol exposure was associated with decreased DMS theta power after premature actions (main effect of reward: F(1,493)=6.67,p=0.01).

All statistical comparisons for OFC theta power in females and males following correct actions were not significant (p values >0.05; Figure 8I). In the DMS, EtOH adolescent females had stronger theta power than adolescent sucrose females following correct actions (Tukey post hoc, p=0.00; Figure 8J). In males, adults had stronger DMS theta power than adolescents following correct actions (main effect of reward: F(1,46)=1.99,p=0.17).

Next, functional connectivity between the OFC and DMS was computed by measuring phase synchrony in the theta frequency band (Figure 8K).

In females, adolescents had stronger OFC-DMS synchrony compared to adults following premature actions (main effect of age: F(1,8699)=298.96,p<2x10^-16^), but in males, OFC-DMS synchrony was stronger in animals who drank EtOH in adolescence compared to sucrose rats (main effect of reward: F(1,5669)=381.5,p<2x10^-16^; Figure 8L).

Compared to adults, adolescent females had weaker OFC-DMS synchrony during correct trials (main effect of age: F(1,1236)=9.58,p=0.002; Figure 8M). In males, OFC-DMS synchrony following correct actions was stronger in animals who drank EtOH in adolescence (main effect of reward: F(1,899)=17.20,p=3.68x10-5; Figure 8M).

We observed that adolescent alcohol’s effects were most pronounced in males following premature actions. Regression models were therefore used to determine which LFP measures were the greatest predictors of premature responses in males. In adolescents, the OFC slope was the greatest predictor of premature actions in sucrose males, while the DMS slope best predicted behavioral performance in EtOH males (Figure 9A). Similarly, in adults, OFC power best predicted sucrose male behavior while DMS slope best predicted EtOH male behavior (Figure 9B).

**Figure 9.**
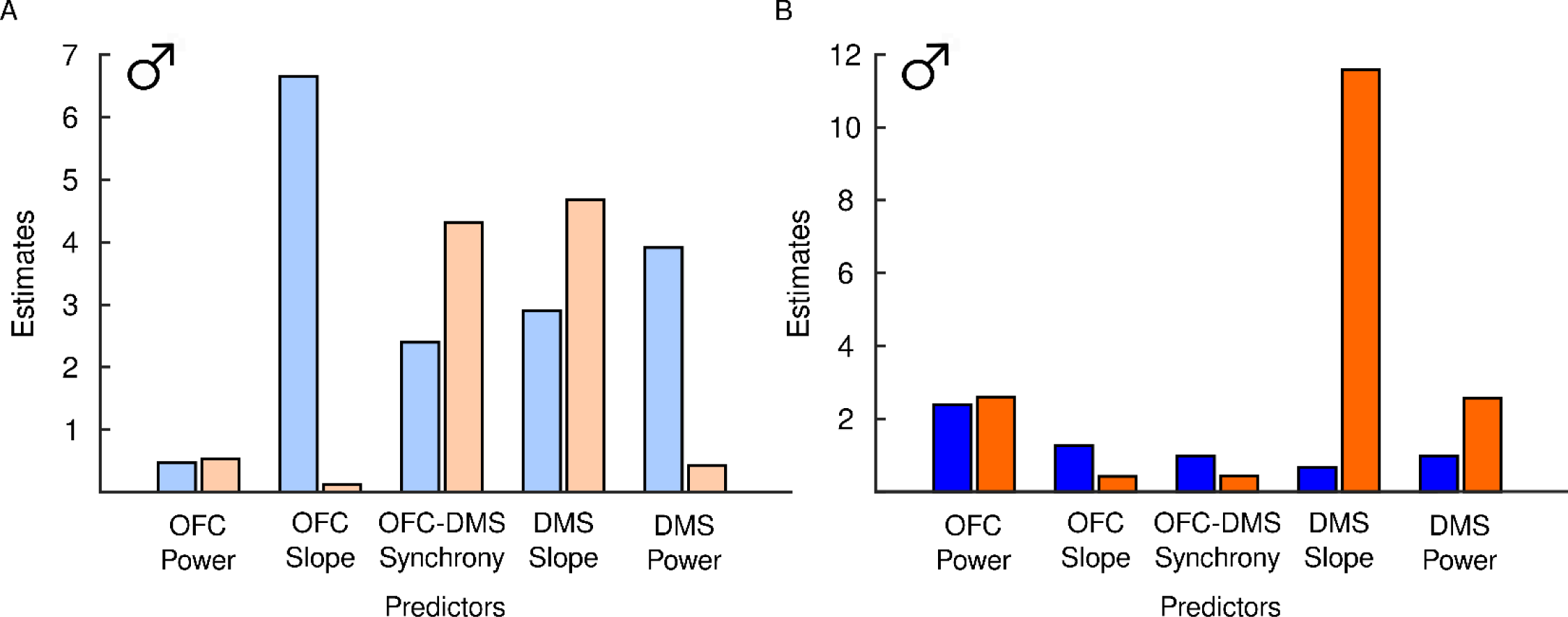
Predictor importance for LFP measures. The ability of each LFP variable to predict number of premature actions was measured in adolescent (A) and adult (B) male rats with a history of EtOH (orange) or sucrose (blue). Higher values indicate greater prediction of behavior. (A) The greatest predictor of premature actions was OFC slope in adolescent sucrose males and DMS slope in adolescent EtOH males. (B) In adult rats, the greatest predictor of premature actions was OFC power in sucrose males and DMS slope in EtOH males.

### Adolescent EtOH exposure promotes increased EtOH intake in adult males but not females

Voluntary EtOH consumption in adulthood was assessed in male and female rats who drank either sucrose or EtOH in adolescence. Overall females drank more EtOH at both the 30 min (main effect of sex: F(1,43)=11.00,p=0.002) and 24-hour time point (main effect of sex: F(1,55)=7.86,p=0.007). Data were therefore divided by sex. EtOH intake after 30 min increased over days in both females (main effect of day: F(6,167)=5.55,p=2.28x10^-5^) and males (main effect of day: F(6,156)=10.05,p=2.25x10^-9^; Figure 10A). Although adolescent alcohol did not significantly impact EtOH intake in either sex (Female: main effect of group: F(1,23)=0.79,p=0.38, Male: main effect of group: F(1,20)=3.39,p=0.08), the males trended towards significance. Female EtOH intake after 24 hrs of access was not influenced by day (main effect of day: F(6,182)=1.81,p=0.10) or adolescent drinking group (main effects of group: F(1,27)=0.07,p=0.80). In contrast, males who experienced adolescent EtOH exposure drank more EtOH in adulthood compared to animals with a history of sucrose (main effect of group: F(1,28)=4.57,p=0.04; Figure 10B).

**Figure 10.**
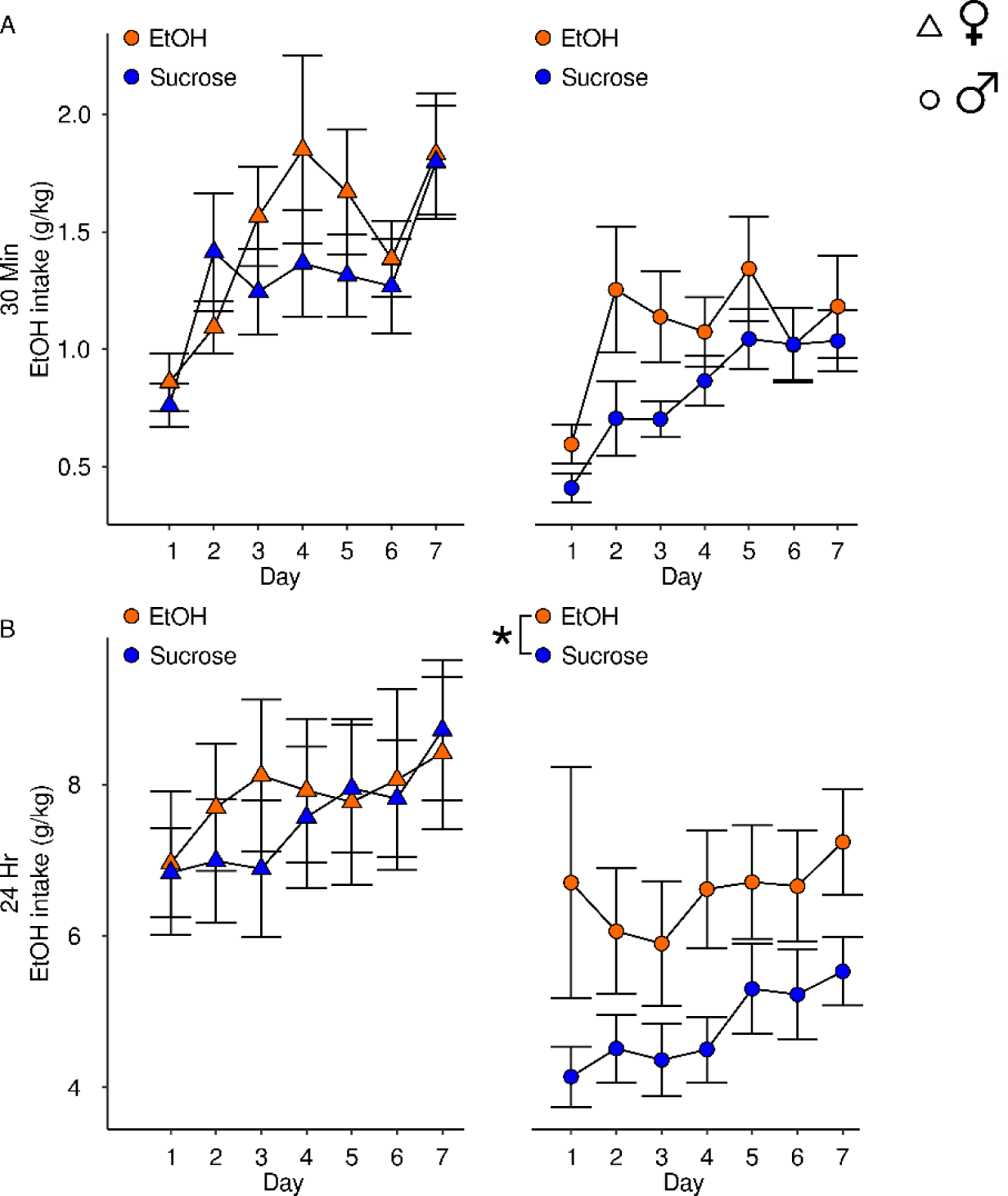
Adult home-cage drinking of EtOH. EtOH intake in female (triangle) and male rats (circle) who drank EtOH (orange) or sucrose (blue) in adolescence after 30 min (A) or 24 hrs access (B) of EtOH. (A) EtOH intake after 30 min increased over days in females and males (B) Female EtOH drinking at the end of 24 hrs was not influenced by adolescent drinking experience. Males who were exposed to EtOH in adolescence drank more EtOH than sucrose exposed males. Data are presented as mean + SEM. *p<0.05 main effect of group

## DISCUSSION

AUD is associated with initiation of drinking during adolescence and with deficits in response inhibition, a behavior mediated in part by OFC-DMS circuitry. Here we investigated the effects of adolescent EtOH exposure on response inhibition while measuring unit and LFP activity in the OFC and DMS in adult and adolescent rodents. Previous work which had reported on adolescent EtOH exposure on response inhibition and physiological correlates in adults (Acheson et al., 2013; Gass et al., 2014; Sanchez-Roige et al., 2014; Passos et al., 2015) had used long EtOH exposure paradigms. Here we used a voluntary (Holgate et al., 2017) and relatively brief drinking paradigm in adolescents. We focused on the OFC and DMS because these regions undergo pronounced development during adolescence and are reported to mediate response inhibition and impulsive choice (Chudasama et al., 2003; Rieger et al., 2003; Eagle et al., 2007; Mar et al., 2011; McCane et al., 2024). Cortical-striatal circuitry is also implicated in in AUD pathology in clinical populations (Cservenka and Nagel, 2012; Courtney et al., 2013; Cservenka et al., 2014). We therefore hypothesized that OFC-DMS activity and connectivity would be vulnerable to the impact of adolescent alcohol drinking and corresponding deficits in response inhibition.

### Adolescent EtOH drinking exerts sex-specific differences in behavior and OFC-DMS activity

Sex-specific effects of EtOH were detected despite the observation that male and female adolescents drank similar levels of EtOH in their home cages. Adolescent EtOH drinking had pronounced effects on males which were unobserved in females. Most notably, adolescent EtOH drinking was associated with increased premature actions, reduced RIRs, faster premature actions, and increased perseverative responding during the ITI, in males but not females. Sex-dependent differences in developmental maturation may explain these differences. Female rats reach puberty around PND 36 while males show signs of sexual maturity completion around PND 40 (Vetter-O’Hagen and Spear, 2012). EtOH exposure which occurs in pre-pubertal to pubertal rats resulted in more deleterious behavioral and physiological effects compared to exposure later in adolescence (Spear, 2015). Additionally, sex differences in gonadal maturation during adolescent alcohol exposure are hypothesized to play a causal role in sex-specific effects of adolescent alcohol exposure on HPA axis-related brain circuitry (Logrip et al., 2013). Thus, because the drinking paradigm utilized was implemented from PND 30-35, males were more likely to be sexually immature which may have resulted in increased vulnerability to the effects of EtOH.

Although adolescent EtOH exerted a weaker influence on female behavior, several physiological differences were observed. In adolescent females, pronounced EtOH-mediated effects were observed in the OFC, but population analyses suggest these effects may be mediated by a relatively small number of units, warranting future exploration. In contrast, adult females who drank EtOH in adolescence show lasting alterations in neural activity. In adulthood, adolescent EtOH exposure caused a suppression in OFC firing rate after premature actions, an increase in OFC firing rate after correct actions, and an attenuated suppression in DMS firing rate following premature actions. The presence of physiological differences in the absence of behavioral differences has two implications. First, the behavioral paradigm may not have been sensitive enough to detect sex-specific strategies mediating behavioral differences. Males and females have been reported to engage in sex-specific strategies when navigating behavioral tasks. For example, females were slower to adapt search strategies, but acquired Pavlovian approach faster than males (Hammerslag and Gulley, 2014; Macht et al., 2020), collectively suggesting that males and females can apply different strategies during cued behaviors. Second, these changes may reflect a change in the way female brains encode premature and correct actions. Adolescent alcohol induced sex-specific changes in physiology but not behavior have been observed previously. Adolescent alcohol drinking had sex-specific effects on molecular markers of serotonergic, dopaminergic, and glutamatergic signaling in adult rats, but resulted in similar recognition memory impairments in both sexes (Marco et al., 2017). Adolescent alcohol exposure was also associated with sex-specific effects on anxiety circuitry and contextual renewal of fear memories (Chandler et al., 2022; Healey et al., 2022). While adolescent alcohol exposure did not influence response inhibition and adult EtOH drinking in females, the data presented here suggest that adolescent alcohol exerts changes on female physiology which may influence behavioral strategies in other tasks. When evaluating sex differences, it is important to assess strategy rather than pure performance metrics (Grissom et al., 2024). Studies which further investigate sex-specific strategies during behavioral performance are therefore necessary to thoroughly understand the impact of adolescent EtOH drinking on female behavior and physiology.

### The effects of moderate EtOH drinking in adolescence parallel chronic EtOH exposure in adults

Adolescent rats consumed moderate amounts of EtOH over days. Despite the short duration of this drinking paradigm and the moderate intake levels, pronounced alterations in physiology and behavior were observed, resembling some of the changes reported after chronic EtOH exposure in adult rodents. For example, chronic alcohol drinking is associated with reduced GABA transmission but increased glutamate transmission in the DMS (Wilcox et al., 2014; Ma et al., 2017; Cuzon Carlson et al., 2018; Renteria et al., 2018; Baltz et al., 2023), reflecting a shift in the excitation/inhibition ratio in favor of excitation. Additionally, the excitation/inhibition balance undergoes profound changes in adolescence (Brenhouse and Andersen, 2011; Spear and Swartzwelder, 2014; Caballero et al., 2021). Our data suggest that even moderate adolescent alcohol use may alter the excitation/inhibition ratio in male rats. We used the aperiodic exponent or power spectrum density slope as an index of excitation-inhibition balance as reported previously (Gao et al., 2017; McCane et al., 2024).

Larger exponents, which are hypothesized to reflect greater inhibition (Gao et al., 2017), were observed in adults compared to adolescents. Moreover, adolescent EtOH exposure produced an apparent, but not statistically significant reduction in slope, reflecting reduced inhibitory transmission in the DMS. Greater striatal inhibition has been linked to impulsive actions in clinical populations. Children with attention deficit hyperactivity disorder exhibit a steeper slope during inhibitory processing compared to controls (Pertermann et al., 2019). Importantly, this difference was ameliorated with the impulsivity treatment methylphenidate (Pertermann et al., 2019), suggesting that increased excitation, indexed by steeper slopes, plays a causal role in impulsive responding. Results from the regression analyses indicated that the DMS slope exerted the greatest influence over premature actions in EtOH-exposed male rats. In contrast, OFC slope and power were the greatest predictors of premature actions in sucrose-exposed adolescent and adult males. Collectively, these data suggest a shift from top-down cortical control over behavior to bottom-up striatal control in male rats with a history of EtOH drinking. Importantly, striatal control over behavior has been hypothesized to lead to impulsive actions (Naaijen et al., 2015) and is a consequence of chronic EtOH exposure (Everitt and Robbins, 2005; Kalivas and Volkow, 2005; DePoy et al., 2013).

### Adolescent EtOH drinking causes a persistent adolescent phenotype

Our data suggest that adolescent alcohol drinking significantly changes OFC-DMS connectivity through adulthood. Adolescents make more premature actions than adults and exhibit an accompanying increase in OFC-DMS connectivity (McCane et al., 2024). We observed that adolescent EtOH drinking is associated with increased premature responding and an accompanying increase in OFC-DMS connectivity in male rats. Increased OFC-striatal connectivity is associated with impulsive and compulsive behaviors (Burguière et al., 2013; Hu et al., 2019). Our data suggest that adolescent alcohol exposure has the propensity to alter brain regions which drive compulsive and impulsive actions, and in this way may promote AUD-pathology in adulthood. Importantly, the “lock-in” of an adolescent phenotype in adult animals who were exposed to alcohol in adolescence has been described previously (Spear and Swartzwelder, 2014; Crews et al., 2016, 2019). However, to our knowledge, this is the first study to show this “lock-in” in cortical-striatal circuits. These data therefore expand upon a rich literature describing adolescent alcohol induced persistent alterations in neurobiology and behavior to show that adolescent alcohol drinking induces persistent alterations to OFC-striatal circuitry maturation.

### Adolescent EtOH drinking increases EtOH intake in male adults

Alcohol drinking in adolescence promotes AUD symptoms in adulthood (Dawes et al., 2000; Crews et al., 2016; Tetteh-Quarshie and Risher, 2023) and is associated with greater likelihood of developing an AUD in adulthood (Buchmann et al., 2009). We therefore investigated the effect of adolescent alcohol drinking on alcohol consumption in adults. We utilized the intermittent access protocol (Simms et al., 2008) which allowed longer access periods of alcohol over two weeks. Adolescent alcohol drinking resulted in increased EtOH consumption in male but not female adults. Sex differences in adult EtOH consumption following adolescent EtOH exposure have been reported on some studies but are absent in others (Robinson et al., 2021). Robinson and colleagues (Robinson et al., 2021) report that several parameters such as drinking model and rodent species are likely to influence the degree that sex influences the impact of adolescent EtOH on adult EtOH consumption. For example, EtOH vapor exposure increased alcohol intake in male but not female mice (Maldonado-Devincci et al., 2022), while a combination of EtOH vapor and voluntary consumption resulted in increased EtOH intake in both male and female rats, with females drinking more than males (Amodeo et al., 2018). Thus, it is unclear whether different drinking models would have yielded sex differences in adult EtOH consumption. Importantly, the age of drinking is another critical parameter, where early adolescent males (PND 30-43) were more vulnerable to the increased EtOH intake in adulthood compared to later adolescent males (PND 45-58) (Alaux-Cantin et al., 2013). Given the different developmental maturation timelines of male and female rats (Walker et al., 2017), it is possible that shifting the period of drinking in females may have resulted in increased EtOH consumption in adulthood.

### Study limitations and future directions

The CRIT paradigm was used to investigate the impact of adolescent alcohol exposure on response inhibition. Importantly, under our current design, we are unable to differentiate between learning and performance as discussed elsewhere (McCane et al., 2024). Thus, it is unclear from our results whether adolescent alcohol produced increased impulsivity or impaired the ability to learn inhibitory responding. It is also possible that EtOH-mediated differences are caused by aberrant encoding of value. There is strong evidence that the OFC encodes information from previous actions to influence future action selection (Cazares et al., 2022).

Moreover, OFC-DMS interactions may play a role in encoding the objective value of a reward (Gore et al., 2023). Therefore, adolescent EtOH drinking may impair value encoding, preventing action-outcome learning during premature and correct actions.

We used a modified drinking paradigm from Holgate and colleagues (2017) to expose adolescent rats to EtOH or sucrose. While most animals drank clinically relevant volumes of EtOH in this paradigm, there was variability in intakes. As such, animals who drank at or below volumes reported in a leak cage were removed from the study. Although the use of high and low drinking animals is a strength of this paradigm, we were underpowered to analyze the data from high and low drinking animals separately. Understanding the neural mechanisms that drive heavy and non-heavy EtOH consumption in animal models is a clinically relevant target that should be addressed in future work.

We used custom-fabricated electrophysiology probes to investigate the neural substrates of response inhibition. While this technique generated several novel findings, it is subject to several shortcomings. We were limited in the number of units collected from each animal, precluding our ability to employ large-scale population analyses. We therefore chose to present overall firing rate, individual unit activity and a distribution of the population-level responses following events of interest. This approach indicated that in some cases, overall responses were largely driven by the activity of small subpopulations. Because of the number of overall units recorded, it is sometimes unclear whether group changes are due to the individual neurons sampled. Future work implementing high-yield techniques should resolve this discrepancy.

## Conclusions

Despite some limitations, data presented here demonstrate that adolescent alcohol drinking alters the development of OFC-striatal circuitry and promotes increased EtOH consumption in adulthood in a sex-specific manner. These data add to a growing line of research investigating the impact of adolescent alcohol exposure and have mechanistic implications for AUD with a developmental etiology.

## Supplemental Figures

**Supplemental Figure 1.**
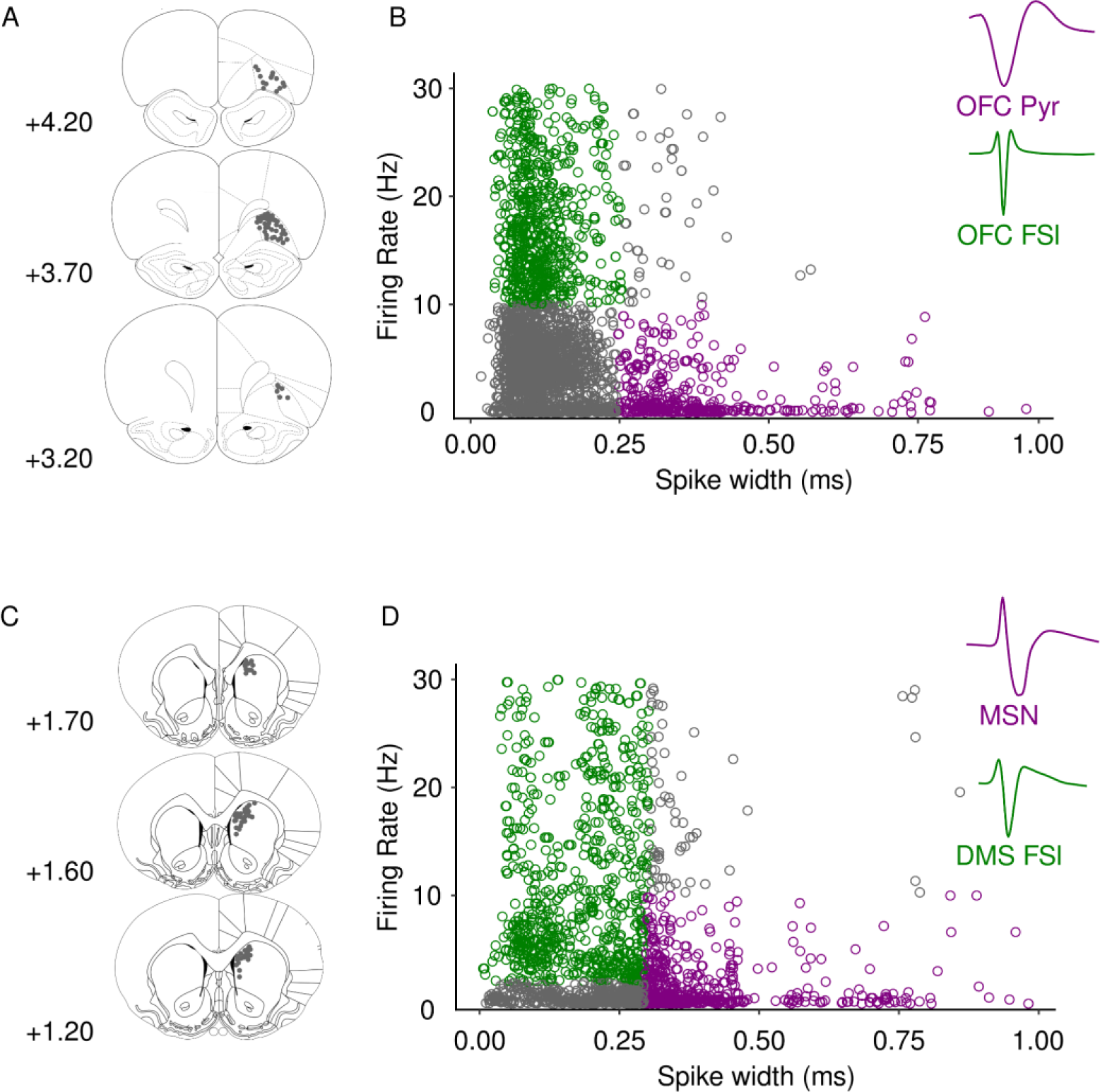
Histology and cell classification. (A) Histological placement of OFC probes. (B) Classification of OFC Pyramidal cells (Pyr; purple), fast spiking interneurons (FSI; green), and representative waveforms. (C) Histological placement of DMS probes. (D) Classification of DMS medium spiny neurons (MSN; purple) and fast spiking interneurons (FSI; green), and representative waveforms.

**Supplemental Figure 2.**
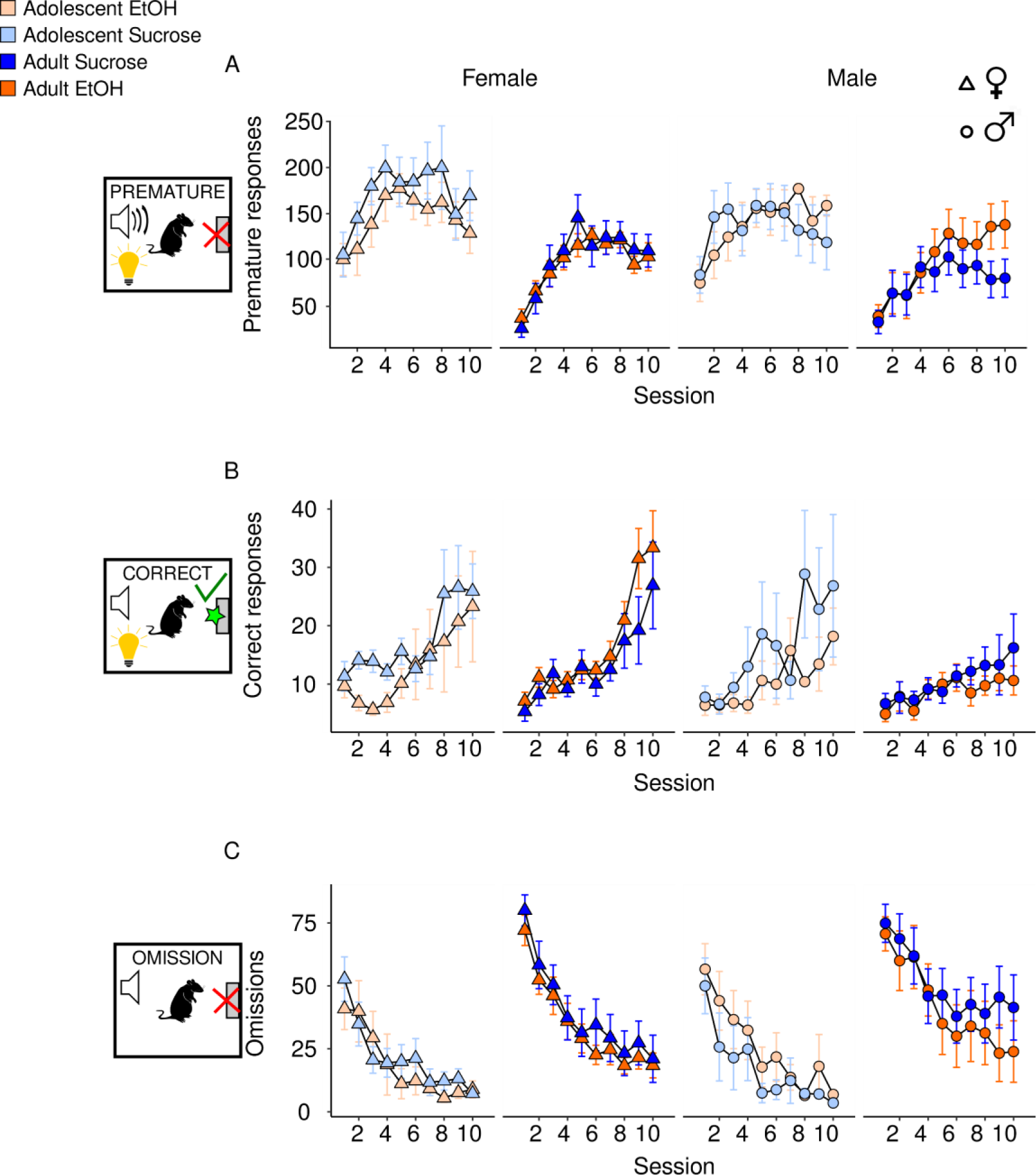
Session by Session CRIT behavioral performance. Behavior in CRIT can be segregated by three distinct trial types: (A) Premature, (B) Correct, and (C) omission. Male (circle) and female (triangle) rats who drank EtOH (orange) or sucrose (blue) performed ten days of CRIT. Lighter shades of each color reflect adolescents and darker shades reflect adults. Bar graphs depict behavioral performance for each group aggregated across the last four sessions of CRIT. Both the number of premature (A) and correct (B) responses increased over session, while the number of omissions decreased for both males and females (C). Data are presented as mean ± SEM.

**Supplemental Figure 3.**
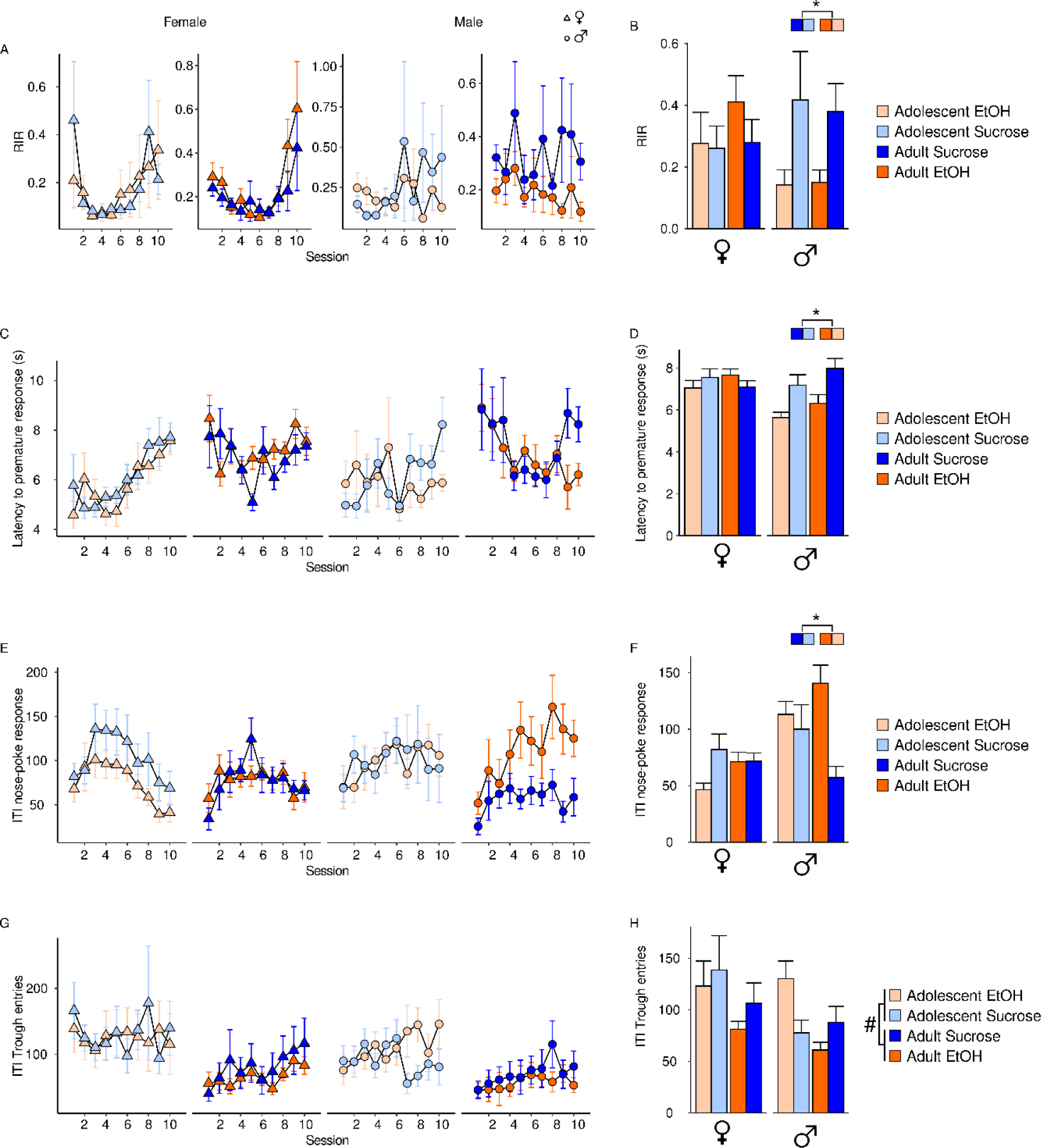
CRIT Behavior. Behavioral measures across session (A,C,E,G) and aggregated across last 4 sessions (B,D,F,H) in female (triangle) and male (circle) rats who consumed EtOH (orange) or sucrose (blue). Adolescents are represented by the lighter shade of each color. (A) Response inhibition ratio (RIR) increased across session in females but not males. (B) RIR over the last four CRIT sessions was reduced in males who drank EtOH in adolescence, but not females. (C) Latency to make a premature response changed across sessions in adolescent and adult females and adult males. (D) Adolescent EtOH exposure was associated with reduced latencies to make a premature response in males but not females. (E) Number of nose-poke responses during the ITI changed across session in females but not males. (F) Adolescent alcohol drinking is associated with increased responding during the ITI in males but not females. (G) Number of trough entries during the ITI changed in female adults only. (H) Adolescents made more ITI trough entries than adults. Data are presented as mean ± SEM. †p<0.05, main effect of sex, #p<0.05 main effect of age, *p<0.05 main effect of group.

**Supplemental Figure 4.**
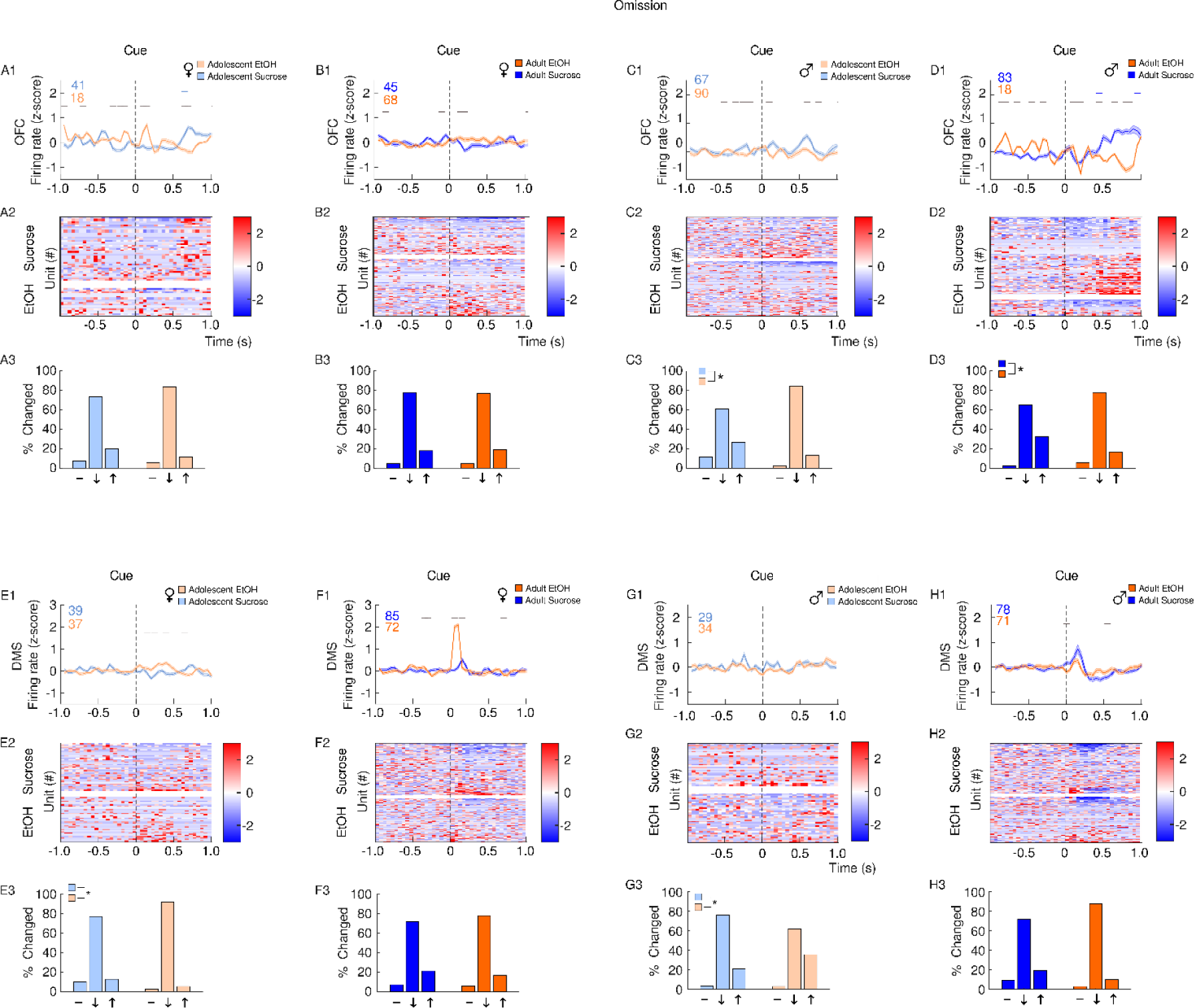
Single unit activity following Omissions. Recordings were performed in the OFC (A-D) and DMS (E-H). Panels depict averaged firing rates (1), heatmaps of individual neural responses (2) and histograms (3) of the distribution of units unchanged (-), inhibited (↓) or excited (↑), 500 ms after cue presentation in male and female rats with a history of EtOH (orange) or sucrose (blue). Adolescents are plotted in the lighter shade of both colors. Events occur at time 0. In adolescent females, OFC firing rate after presentation of cues which preceded omission trials showed no apparent pattern in the overall firing rate (A1), population responses (A2) or the distribution of neural responses (A3). In adult females, OFC firing rate after presentation of cues which preceded omission trials showed no apparent pattern in the overall firing rate (B1), population responses (B2) or the distribution of neural responses (B3). In adolescent males, OFC firing rate after presentation of cues which preceded omission trials showed no apparent pattern in the overall firing rate (C1), but in the EtOH group there is an increased subpopulation of neurons that become inhibited (C2) to a greater extent than in the sucrose group (C3).In adult males, OFC firing rate after presentation of cues which preceded omission trials shows a transient increase in sucrose but not EtOH rats (D1), which is apparent in the population level response (D2). Moreover, a greater proportion of OFC neurons in the sucrose group become excited following cue presentation compared to the EtOH group (D3). In adolescent females, DMS firing rate after presentation of cues which preceded omission trials was greater in EtOH rats compared to sucrose rats (E1). While both groups had a subpopulation of excited neurons (E2), a grater proportion of DMS neurons in the EtOH group were inhibited compared to the sucrose group (E3). In adult females, DMS firing rate after presentation of cues which preceded omission trials was larger in EtOH experienced rats (F1), but both population responses (F2) and the distribution of responses were statistically similar between groups (F3). In adolescent males, DMS firing rate after presentation of cues which preceded omission trials showed no apparent pattern in the overall firing rate (G1), or population responses (A2), but sucrose animals had a larger proportion of neurons inhibited compared to EtOH rats (G3). In adult males, DMS firing rate after presentation of cues which preceded omission trials was slightly increased in sucrose rats compared to EtOH rats (H1). A subpopulation of excited and inhibited neurons was observed in both sucrose and EtOH rats (H2), but the distribution of responses was statistically similar between groups (H3). Data are presented as mean ± SEM. Colored numbers reflect the number of units recorded in each group. Colored significant bars represent Tukey HSD post hoc testing (p<0.05), compared to baseline firing rate for each group (EtOH or sucrose). Black significant bars reflect permutation testing between reward groups, p<0.05. *p<0.05 group difference, Chi-squared test

